# SEHI-PPI: An End-to-End Sampling-Enhanced Human-Influenza Protein-Protein Interaction Prediction Framework with Double-View Learning

**DOI:** 10.1101/2025.03.07.642105

**Authors:** Qiang Yang, Xiao Fan, Haiqing Zhao, Zhe Ma, Megan Stanifer, Jiang Bian, Marco Salemi, Rui Yin

## Abstract

Influenza continues to pose significant global health threats, hijacking host cellular machinery through protein-protein interactions (PPIs), which are fundamental to viral entry, replication, immune evasion, and transmission. Yet, our understanding of these host-virus PPIs remains incomplete due to the vast diversity of viral proteins, their rapid mutation rates, and the limited availability of experimentally validated interaction data. Additionally, existing computational methods often struggle due to the limited availability of high-quality samples and their inability to effectively model the complex nature of host-virus interactions. To address these challenges, we present SEHI-PPI, an end-to-end framework for human-influenza PPI prediction. SEHI-PPI integrates a double-view deep learning architecture that captures both global and local sequence features, coupled with a novel adaptive negative sampling strategy to generate reliable and high-quality negative samples. Our method outperforms multiple benchmarks, including state-of-the-art large language models, achieving a superior performance in sensitivity (0.986) and AUROC (0.987). Notably, in a stringent test involving entirely unseen human and influenza protein families, SEHI-PPI maintains strong performance with an AUROC of 0.837. The model also demonstrates high generalizability across other human-virus PPI datasets, with an average sensitivity of 0.929 and AUROC of 0.928. Furthermore, AlphaFold3-guided case studies reveal that viral proteins predicted to target the same human protein cluster together structurally and functionally, underscoring the biological relevance of our predictions. These discoveries demonstrate the reliability of our SEHI-PPI framework in uncovering biologically meaningful host-virus interactions and potential therapeutic targets.

## Introduction

Influenza remains a significant global health concern, causing annual epidemics and occasional pandemics that result in substantial morbidity and mortality worldwide^1^. According to the CDC, influenza caused an estimated 9.3–41 million illnesses, 100,000–710,000 hospitalizations, and up to 51,000 deaths annually in the United States between 2010 and 2023^2^. In the recent 2024-2025 flu season, Flu cases have surged to the highest levels in 15 years since the swine flu pandemic in late 2009, and influenza activity remains elevated and continues to increase across the country. Concurrently, there has been a notable rise in avian influenza (H5N1), with the virus spreading to mammals, raising concerns about its potential to infect humans^3^. Such cross-species could lead to immune escape, resulting in large-scale infections and posing a severe threat to global health. The influenza virus consists of 8 different segments (proteins), where HA (hemagglutinin) is the most critical one that enables the viral particle to infect host cells. The HA protein attaches to the host cell by binding to sialic acid receptors through the host’s receptor proteins^4^. Within the acidic environment of the endosome, HA undergoes a conformational change. This change exposes the fusion peptide, which facilitates the fusion of the viral envelope with the endosomal membrane, ultimately releasing the viral genetic material into the cytoplasm for replication^5,6^. Once inside the host cell, the influenza virus hijacks cellular machinery to replicate and produce viral components, disrupting normal cellular functions. Apart from the HA protein, other viral segments (proteins) also play critical roles during viral infection. For instance, the non-structural protein 1 (NS1) suppresses host immune responses by inhibiting interferon production and manipulating host RNA processes^7^, while the Matrix Protein 2 (M2) alters ion balance to facilitate uncoating and assembly^8^. The infection culminates with the assembly of new virions, aided by the neuraminidase (NA) protein, which cleaves sialic acid residues to release the virions and prevent aggregation^9^. These intricate processes highlight the virus’s ability to exploit host cellular mechanisms for replication and propagation, significantly enhancing its infection efficiency.

Understanding host-virus protein-protein interactions (PPIs) is fundamental for unraveling the mechanisms of viral replication and pathogenesis during human infection^10^, particularly amid rising concerns over the spread of highly pathogenic avian influenza H5N1. In early 2025, the detection of H5N1 in diverse species, including dairy cattle in the United States and, for the first time, sheep in the United Kingdom, has raised alarm over its potential to cross species barriers and infect humans^11,12^. Host-virus PPIs represent the molecular interface at which viruses exploit host cellular machinery to promote their own replication, evade immune responses, and ultimately cause disease. Biological experiments, including yeast two-hybrid^13^ (Y2H), mass spectroscopy^14,15^ (MS), co-immunoprecipitation^16^, and tandem affinity purification^17^ (TAP), have been employed to identify human-virus PPIs. However, these methods are costly, time-consuming, and prone to high false-positive or false-negative rates^18^. Moreover, given the immense combinatorial space of possible protein interactions, experimentally mapping all potential host-virus PPIs remains infeasible. Consequently, existing datasets remain sparse and incomplete, especially for newly emerging or less-studied viruses like H5N1. To complement experimental approaches, computational methods have emerged as essential tools for PPI prediction. Early computational methods focused on leveraging the genomic and biological context of proteins to generate interaction networks and identify functional linkages^19^. Though these approaches are feasible and sound, they heavily rely on indirect associations, such as genetic linkage and evolutionary conservation^20^. Subsequent methods combined classical machine learning algorithms, including support vector machines, decision trees, and XGBoost, with manually engineered features derived from protein sequences for PPI prediction^21^. Such approaches, albeit effective, cannot automatically extract deep-level features of PPI from the original sequences or structures of proteins, which hampers further improvements in predictive performance and suffers from poor generalizability.

In recent years, deep learning-based methods have revolutionized host-virus PPI prediction. In detail, these methods can be classified into several categories based on: (1) graph neural networks (GNNs)^22,23^, (2) convolutional neural networks (CNNs)^24,25^, (3) recurrent neural networks (RNNs)^26,27^, and (4) hybrid architecture^28,29^, etc. Most of these methods extract features directly from source data (i.e., protein sequences and 3D structures) to identify complex interactions among proteins (e.g., the relationships between sequence motifs, structural properties, and physicochemical attributes) for PPI prediction. For structure-based methods, Baranwal et al. proposed to use a multi-layer graph attention network to predict PPIs solely from 3D structural information, i.e., atom positions of folded protein globules^22^. For feature-based methods, Gao et al. integrated various feature types, including the auto covariance descriptor, multivariate mutual information, and conjoint triad, and employed a combination of residual convolutional neural network, fully connected network, LightGBM, and XGBoost for PPIs prediction^30^. For sequence-based approaches, Yang et al. designed a deep learning framework by representing interacting protein sequences with a pre-acquired protein sequence profile module followed by a Siamese CNN and a multi-layer perceptron (MLP) module^18^. Soleymani et al. put forward an autoencoder architecture to encode protein sequences into a lower-dimensional vector while preserving its underlying sequence attributes^31^. Nambiar et al. presented a Transformer neural network to pre-train task-agnostic sequence representations and fine-tuned it for PPI prediction^32^. For hybrid approach methods, Yang et al. established a multi-modal embedding feature fusion-based LightGBM method to predict human-herpesvirus PPIs^33^. Despite these advancements, existing methods face several limitations: 1) Structure-based methods mainly depend on high-resolution protein structures, whereas most of the protein structures are missing, limiting their utility and applicability^34^. 2) Current approaches often fail to capture the intricate and dynamic nature of host-virus PPI interactions, leading to suboptimal predictive performance and poor generalization, particularly to the interactions caused by emerging new viral strains. 3) There is no consensus or ground truth on the generation of negative samples (i.e., non-interaction PPI pairs) and random sampling is currently the most common strategy used, while this strategy often results in low-quality samples for modeling^35^.

To overcome the above discussed technical challenges, in this study, we propose an end-to-end **<u>s</u>**ampling-**<u>e</u>**nhanced **<u>h</u>**uman-**i**nfluenza **<u>PPI</u>** prediction framework, named **SEHI-PPI**, whose overall architecture is shown in **Figure 1**. In SEHI-PPI, we proposed an adaptive negative sampling strategy through *K*-mer sub-sequences to generate non-interacting PPI pairs. To better capture the intricate and dynamic nature of viral sequences, we came up with a new double-view feature extraction module. In this module, sequences are treated as graph-like, where their global-view features are extracted by recurrent neural networks to represent the overall patterns, while their local-view features are derived from graph neural networks to represent the local sequence specificity. To assess its generalizability, we further evaluated SEHI-PPI in two key scenarios: (1) predicting PPIs for novel protein families unseen during training, and (2) fine-tuning SEHI-PPI for human-virus PPI prediction beyond influenza. Additionally, we conducted case studies on two human proteins: HSPA1B (heat shock protein family A [Hsp70] member 1B) and RTRAF (RNA transcription, translation, and transport factor). In both cases, we observed a consistent pattern between sequence clustering of their viral protein interactors and structural alignments predicted by AlphaFold 3. Overall, SEHI-PPI introduces a novel negative sampling strategy and an enhanced feature extraction module, achieving superior performance in human-virus PPI prediction. Its strong transferability suggests broad applicability for studying other host-virus interactions beyond influenza. SEHI-PPI has the potential to serve as a surveillance tool, enhancing preparedness for emerging viral outbreaks by facilitating early-stage detection of viral pathogenesis and informing therapeutic and preventive strategies. The main contributions of this work are summarized below:

**Figure 1.**
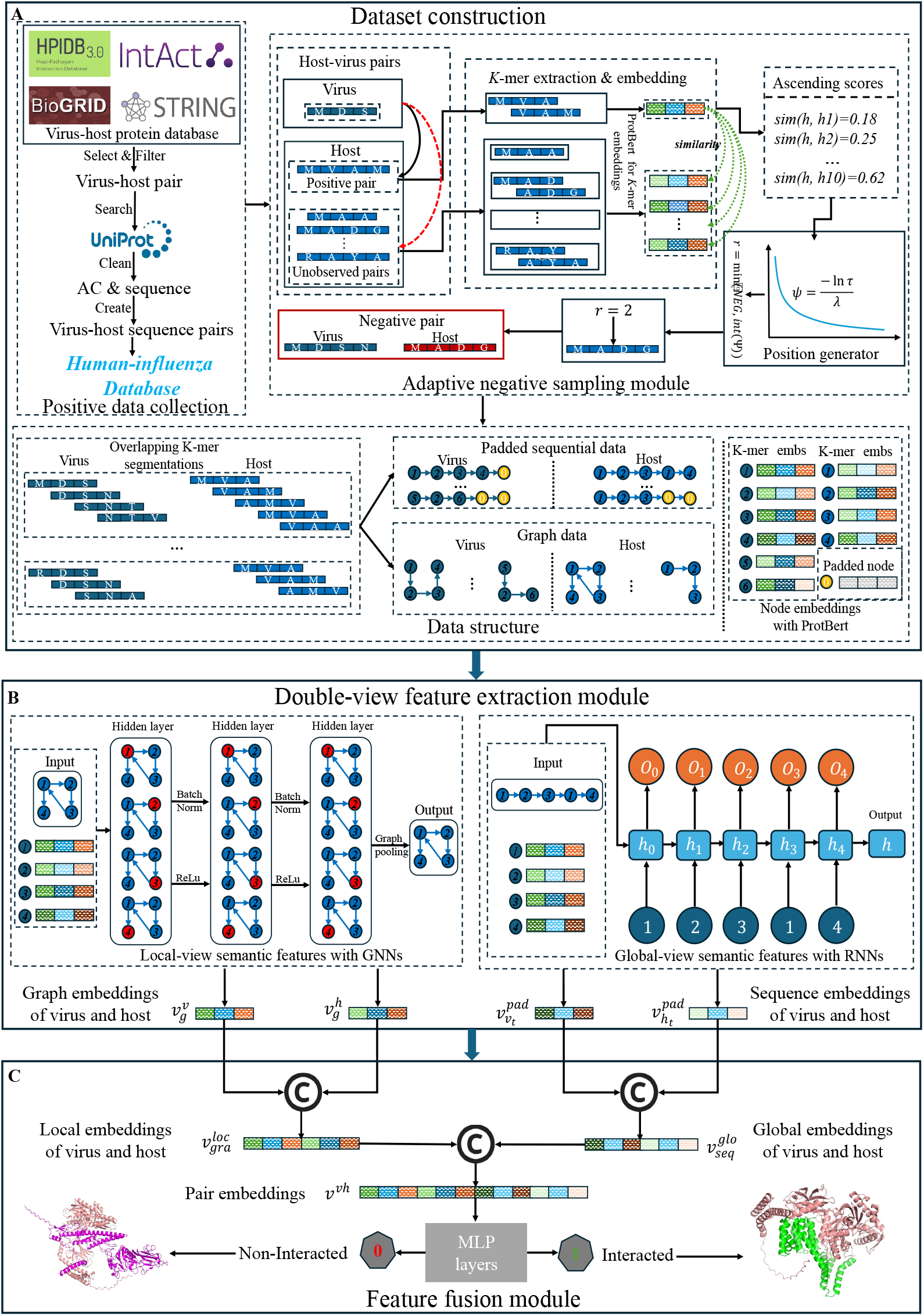
The workflow of our SEHI-PPI model. (A) Data construction involves creating a dataset that combines collected positive data with generated negative data and building sequential and graph data using overlapping *K*-mer segmentations. (B) The double-view feature extraction module is designed to extract the local and global features with GIN and LSTM, respectively. (C) The feature fusion layer is developed to learn the human-influenza pair embeddings, which further are fed into MPL layers for classification.

- We developed an end-to-end generalizable human-influenza PPI prediction framework based on the combined global and local features.
- We proposed a novel adaptive negative sampling strategy for high-quality negative sample generation of human-influenza PPI pairs.
- Extensive experiments have demonstrated the superior performance of our proposed model for human-influenza PPI prediction with comprehensive benchmark comparison.
- We validated the generalization capabilities of our model by applying it to protein-protein pairs, including human and viral protein families not involved in the training process, and to other types of human-virus PPI prediction, demonstrating its adaptability across diverse host-virus interactions.

## Results

### An end-to-end generalized framework with sampling-enhanced double-view learning

#### Overall architecture

In this work, we developed SEHI-PPI, an end-to-end deep learning framework (**Figure 1**), to identify human-influenza PPI incorporating a novel double-view deep learning method and an adaptive negative sampling strategy. SEHI-PPI consisted of three primary modules: dataset construction module, double-view feature extraction module, and feature fusion module for human-influenza PPI prediction. The dataset construction module collected positive PPI pairs from public databases mapped them to UniProtKB sequences and generated high-quality negative (non-interactive) protein-protein pairs using the proposed adaptive negative sampling strategy (**Figure 1A**). Protein sequences were then segmented into *K*-mers and encoded using pre-trained protein language models (PLMs) to create sequential and graph-based representations. The double-view feature extraction module extracted the global and local features with RNNs (e.g., long short-term memory [LSTM]) and GNNs (e.g., graph isomorphism network [GIN]) (**Figure 1B**). The global-view features of protein sequences captured overall patterns and characteristics defining the protein’s function, while local-view features focused on specific sequence elements crucial for precise interactions and activities. Finally, the feature fusion module concatenated global and local embeddings of human and influenza proteins, forming pairwise representations, which were processed through MLP layers to predict human-influenza PPIs (**Figure 1C**). In the following subsections, we provide a detailed explanation of each step. Comprehensive technical details are available in the Methods section and Supplemental Information (SI).

### Performance evaluation of SEHI-PPI on human-influenza PPI dataset

#### Description of datasets

The collected positive dataset comprises 15,889 human-influenza PPI pairs, including twelve different influenza proteins, including non-structural protein 1 (NS1), nucleoprotein (NP), polymerase basic protein 2 (PB2), matrix protein 1 (M1), nuclear export protein (NEP), polymerase acidic protein (PA), hemagglutinin (HA), RNA-directed RNA polymerase catalytic subunit (PB1), neuraminidase (NA), polymerase basic 1-F2 protein (PB1-F2), matrix protein 2 (M2), polymerase acidic X protein (PA-X) and 4277 distinct families of human proteins. The protein distributions of influenza and humans (top 50) are shown in Supplementary Fig. 2. The statistics information of the dataset is shown in Supplementary Table S1. Additionally, we used our proposed adaptive negative sampling strategy to generate an equal number of non-interactive human-influenza protein-protein pairs for modeling. Also, to evaluate the generalizability of our proposed framework, we also collected interactive protein-protein pairs of other human-virus types following the same procedures, including Zika virus (Zika, #4,809), Murid herpesvirus 1 (MHV1, #13,160), Human herpesvirus 1 (HHV1, #23,486), Zaire ebolavirus (Ebola, #2,360), severe acute respiratory syndrome coronavirus (SARS-CoV, #12,334), Beta papillomavirus (BPPLM, #6,521), and Human herpesvirus 8 (HHV8, #11,906).

**Figure 2.**
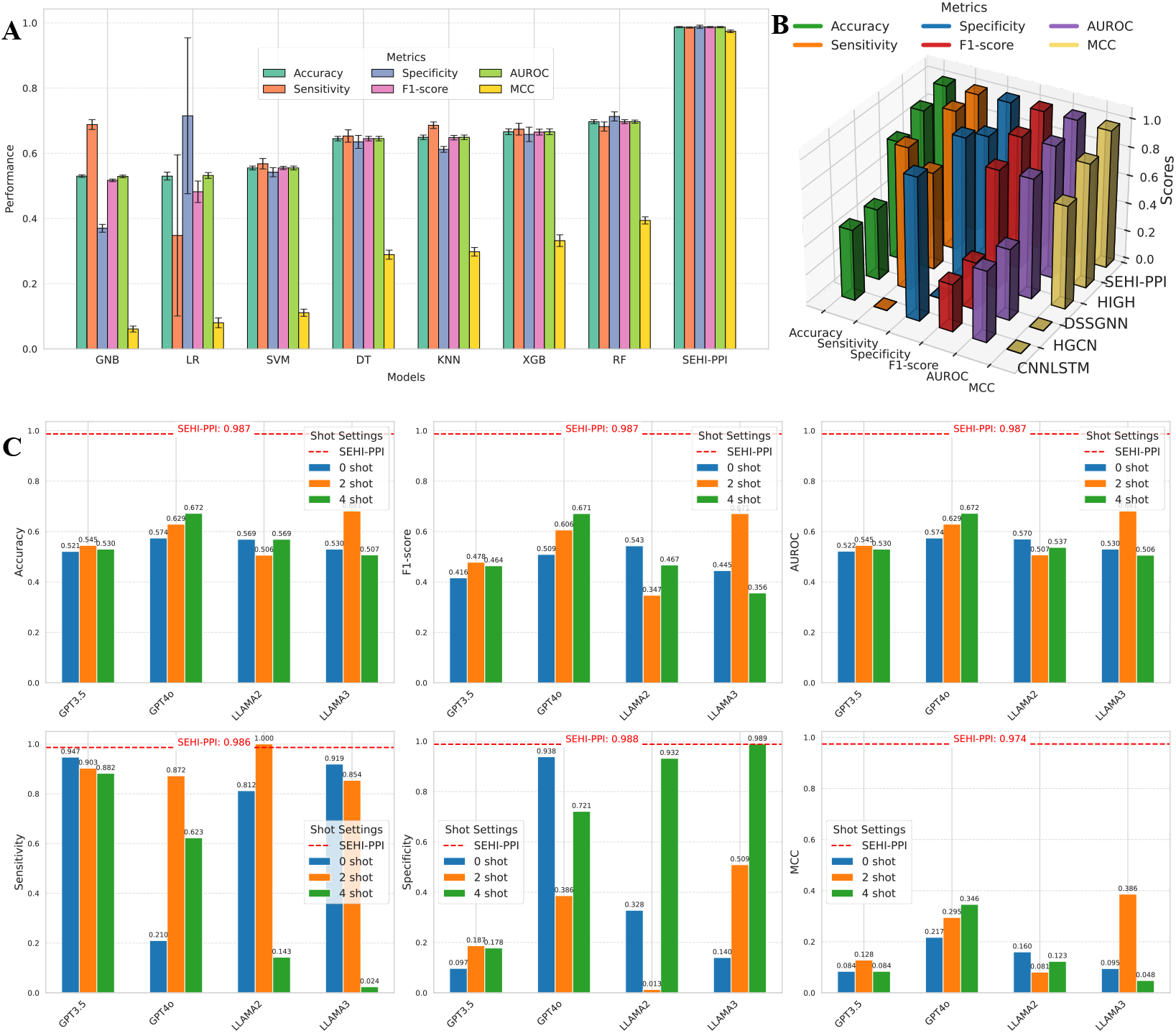
Performance comparison with existing three categorical methods for human-influenza PPI prediction. (A) Performance comparison with traditional machine learning methods (mean ± standard deviation), including Gaussian Naive Bayes, logistic regression, and support vector machine, decision tree, K-nearest neighbor, XGBoost, and random forest. (B) Performance comparison with existing state-of-the-art models, containing CNNLSTM, HGCN, DSSGNN, and HIGH with our SEHI-PPI framework (LSTM+GIN). (C) Performance comparison of different LLMs, involving GPT-3.5, GPT-4, LLAMA2, and LLAMA3 across different shot settings with our SEHI-PPI framework (LSTM+GIN).

#### Performance of SEHI-PPI using different sequence- and graph-based architectures

For the global-view feature extraction, we implemented RNNs with LSTM^27^, and gated recurrent unit (GRU)^36^, while for the local-view feature extraction, we employed GNNs with GraphSAGE^37^, graph convolution network (GCN)^38^, graph attention network (GAT)^39^, principal neighborhood aggregation network (PNA)^40^, and simple graph convolution network (SGC)^41^. Further methodological details were provided in Supplementary Note 1. **Table 1** presents the performance of different combinations of RNNs and GNNs for human-influenza PPI prediction on a testing set using global and local-view features. According to the results, when applying the same RNN architecture, i.e., LSTM, SGC achieves better performance, with the same 0.989 ± 0.001 in accuracy, F1-score, and AUROC, 0.980 ± 0.003 in sensitivity, 0.978 ± 0.001 in specificity, 0.978 ± 0.002 in MCC followed by GIN, GCN, PNA, and GraphSAGE. It can be reasonably inferred that the strengths of SGC streamline graph convolution operations, which effectively model sequential dependencies. GAT has a slightly lower performance with an accuracy of 0.973 ± 0.010, F1-score of 0.973 ± 0.010, and AUROC of 0.973 ± 0.010 while leveraging attention mechanisms for dynamic weighting. This is probably due to the increased model complexity that leads to higher variability. For GRU-based models, GIN and SGC achieve almost the same averaged performance over five-fold runs, however, SGC exhibits smaller variability, demonstrating its greater stability and consistent learning process across the repeated experiments. In addition, when utilizing the same GNN models, i.e., GIN and SGC, we find that LSTM and GRU demonstrate similar performance. This can be attributed to their shared ability to handle long-term dependencies effectively through gating mechanisms, which minimizes the impact of their architectural differences for sequence data. To summarize, SGC consistently outperforms other GNNs across both LSTM- and GRU-based architectures, demonstrating its superior ability to capture sequential dependencies. Meanwhile, LSTM and GRU exhibit comparable effectiveness, further validating their suitability for sequence-driven PPI prediction.

**Table 1.**
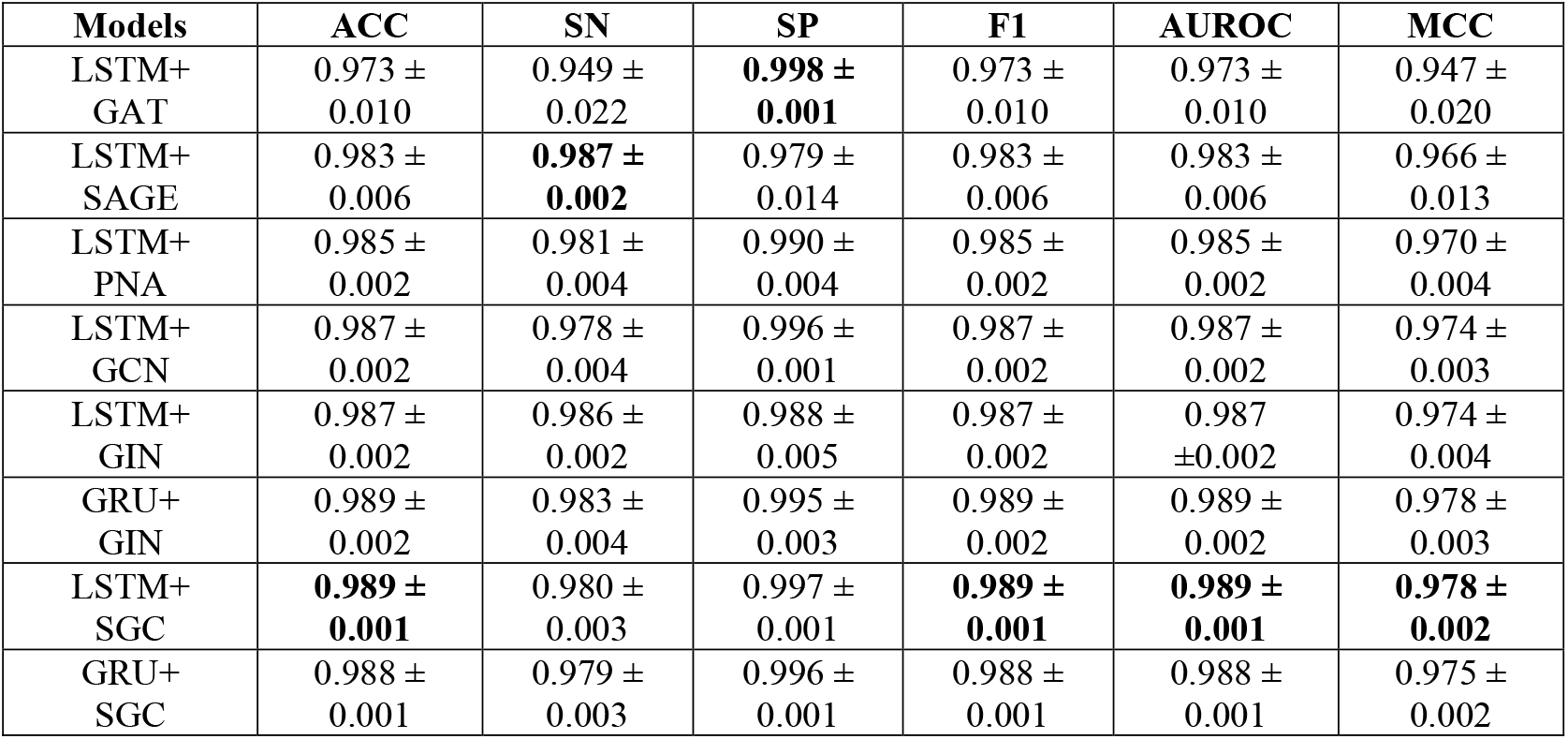
Performance comparison of SEHI-PPI with different GNN and RNN models (mean ± standard deviation). ACC: accuracy; F1: F1-score; SN: sensitivity; SP: specificity; MCC: Matthew’s correlation coefficient.

#### Comparison with traditional machine learning methods

To evaluate SEHI-PPI’s predictive capabilities, we compared it against traditional machine learning models. As shown in **Figure 2**A, among the baseline methods, Gaussian Naive Bayes (GNB), logistic regression (LR), and support vector machine (SVM) exhibit the lowest performance across nearly all metrics. Specifically, GNB achieves an accuracy of 0.530 ± 0.004, F1-score of 0.517 ± 0.004, and AUROC of 0.529 ± 0.004, while SVM shows similar limitations with an accuracy of 0.555 ± 0.012, F1-score of 0.555 ± 0.033, and AUROC of 0.555 ± 0.009. These results reflect their inability to effectively process high-dimensional features (i.e., 1024-dimensional embeddings). While decision tree (DT) and K-nearest neighbor (KNN) classifiers show moderate improvement, they fail to capture complex feature relationships, showing 0.645 ± 0.007 in both accuracy and AUROC. Some other more advanced methods, like XGBoost (XGB) and random forest (RF), deliver better results as seen in **Figure 2**A. In contrast, our proposed SEHI-PPI model, with LSTM+GIN architecture, significantly outperforms all baseline methods, achieving an accuracy of 0.987 ± 0.002, F1-score of 0.987 ± 0.002, and AUROC of 0.987 ± 0.002, with near-perfect sensitivity (sensitivity: 0.986 ± 0.002) and specificity (specificity: 0.988 ± 0.005). These results suggest approximately a 20% improvement over RF, the best-performing baseline model, highlighting the effectiveness of SEHI-PPI in capturing both global sequential and local graph-based features for human-influenza PPI predictions.

#### Comparison with existing PPI prediction methods

We further benchmarked SEHI-PPI implemented by the architecture LSTM+GIN against several existing cutting-edge PPI prediction methods, including CNNLSTM^42^, HGCN^43^, DSSGNN^44^, and HIGH^45^. The results can be found in **Figure 2**B (3D bar chart for performance comparison across models), where CNNLSTM and HGCN exhibit limited predictive capabilities, with both 0.500 in AUROC, while CNNLSTM shows minimum sensitivity (0.001), whereas HGCN indicates perfect sensitivity (0.999). DSSGNN with an AUROC of 0.844, performs moderately well due to its dual-level GNN architecture that integrates protein- and interaction-level features. However, its high specificity (0.998) and lower sensitivity (0.690) also indicate a bias toward negative predictions, showing room for improvement in balancing predictions. HIGH achieves much better results, with an AUROC of 0.936, sensitivity of 0.994, and specificity of 0.878, benefiting from its hierarchical graph structure which may have captured both macro-level PPI relationships and micro-level protein features. Nevertheless, none of these methods compete against our proposed SEHI-PPI, which achieves the best overall performance, owing to its ability to capture both global and local features through its double-view framework. This framework enables a more nuanced understanding of virus-host sequence interactions, positioning SEHI-PPI as a highly effective tool for human-influenza PPI prediction.

#### Comparison with large language models (LLMs)

We also assessed the predictive capability of our framework by comparing it with LLMs, including GPT-3.5^46^, GPT-4^47^, LLAMA2^48^, and LLAMA3^49^ under the setting of zero-shot, two-shot, and four-shot. The designed prompts for zero-shot and few-shot learning are presented in Supplementary Table 2. As shown in **Figure 2**C, the SEHI-PPI framework consistently outperforms almost all LLMs across evaluation metrics. Each subplot features a dashed red line representing SEHI-PPI performance, which exceeds the bars corresponding to LLM results for all shot settings (i.e., 0-shot, 2-shot, and 4-shot). For instance, SEHI-PPI achieves an impressive 0.987 in accuracy, significantly surpassing the best LLM GPT-4 (4-shot, accuracy = 0.672). Notably, while LLAMA2 (2-shot) reaches a perfect sensitivity of 1.000, SEHI-PPI exhibits a competitive performance (0.986). These results indicate that although LLMs benefit from additional labeled examples, their performance remains substantially below that of SEHI-PPI. The consistent superiority of SEHI-PPI, regardless of shot configuration, highlights its ability to capture the intricate semantic and structural relationships inherent in host-virus PPIs, setting a formidable benchmark that LLMs have yet to reach in this domain.

### Ablation studies of SEHI-PPI

#### Adaptive negative sampling promoting the performance of SEHI-PPI

We compared our proposed Adaptive Negative Sampling (ANS) with Random Negative Sampling (RNS) and Statistical Negative Sampling (SNS) under identical settings to assess its contribution to model performance. Each method was first applied to generate negative samples, followed by constructing training, validation, and testing datasets in an 8:1:1 ratio. To ensure a fair comparison, we merged the testing datasets generated by all three methods into a final unified testing dataset, ensuring that the same testing data was used across all sampling strategies for an unbiased performance evaluation. More details about these negative sampling methods can be found in Supplementary Note 2. As summarized in **Figure 3**A, RNS exhibits the lowest performance across all metrics, including accuracy (0.540 ± 0.021), F1-score (0.429 ± 0.087), and AUROC (0.530 ± 0.030), coupled with high variability. This inconsistency reflects the noise introduced during training due to the random selection of negative samples, which lacks informative features. SNS outperforms RNS, displaying higher accuracy (0.677 ± 0.005), F1-score (0.651 ± 0.004), and AUROC (0.681 ± 0.005). These improvements are attributed to the incorporation of *K*-mer overlap analysis, which provides more biologically informative samples, reducing noise to some extent. However, our designed ANS strategy exceeds both approaches, achieving the highest accuracy (0.711 ± 0.001), F1-score (0.693 ± 0.003), and AUROC (0.721 ± 0.004) with reduced standard deviation, indicating superior consistency and lower variance. These results demonstrate two critical advantages of ANS: (1) its ability to optimize model training by adaptively selecting negative samples based on *K*-mer-based sequence similarity and the designed position generator, and (2) its contribution to model generalization, enabling reliable predictions on unknown data for PPI prediction.

**Figure 3.**
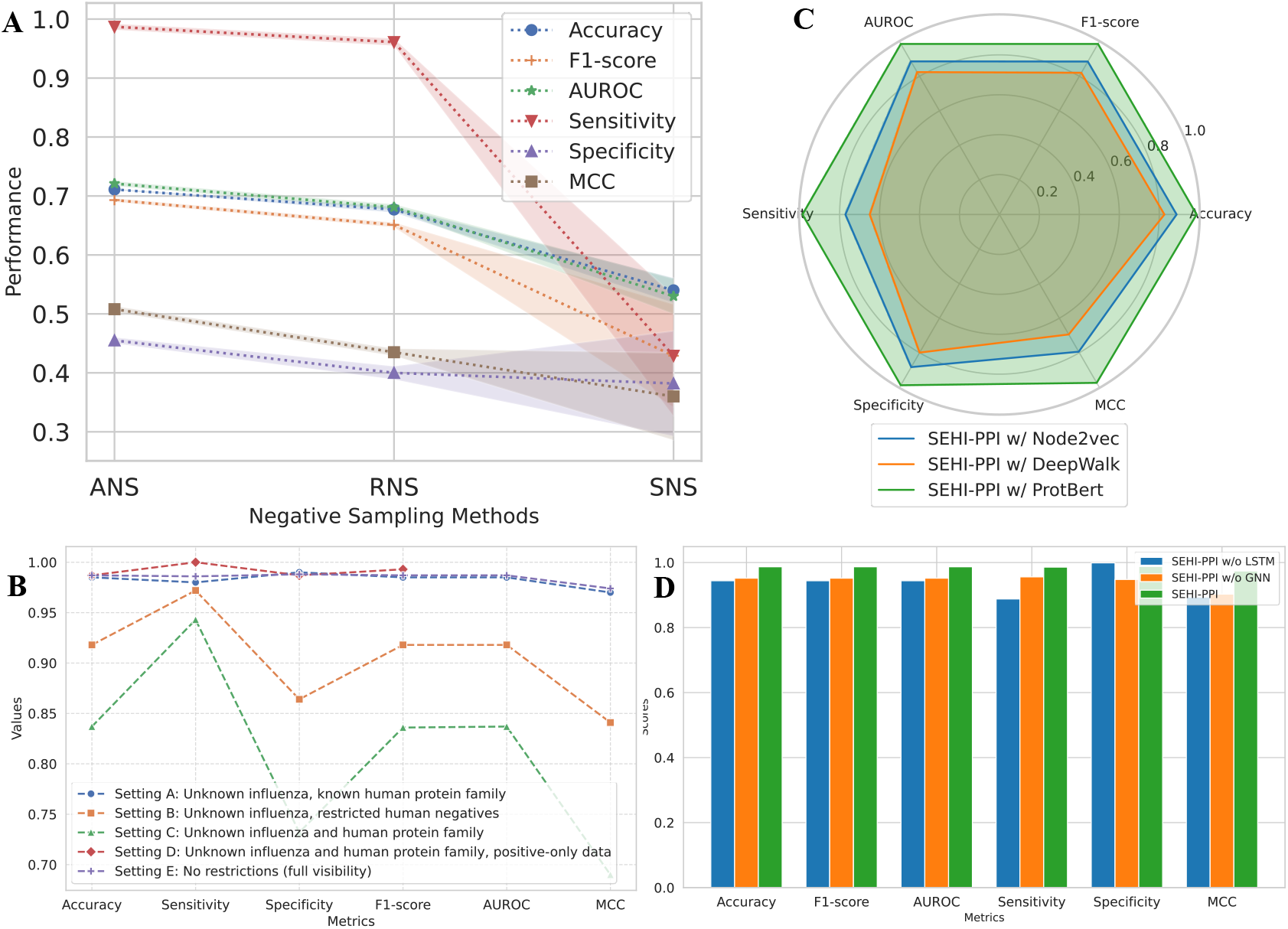
Ablation studies of our proposed SEHI-PPI model. (A) Performance comparison of different sampling strategies. The lines represent mean performance metrics while the shaded regions indicate the variance. Larger shaded areas imply higher variability in performance. (B) Generatability analysis of SEHI-PPI under different human and influenza protein visibility (RE: Recall and PR: Precision). (C) Influence of different *K*-mer-based segmentation embedding methods in SEHI-PPI model. (D) Analysis of global and local feature extraction contributions on SEHI-PPI model performance.

#### Applying SEHI-PPI to novel human and influenza proteins for PPI prediction

We tested SEHI-PPI’s generalizability capability by applying it to predicting human-influenza PPI, of which the protein families were not known by our model. We divided the influenza proteins into training, validation, and testing groups, where the training and validation proteins consist of PB2, M1, NEP, PA, HA, PB1, NA, PB1-F2, M2, and PA-X, and the testing proteins involve NS1 and NP. This setup ensures that viral proteins in the testing data do not appear during training, allowing for a rigorous evaluation of SEHI-PPI’s generalization ability. To perform a comprehensive evaluation, we defined five different situations, represented as Setting A to E, respectively. In Setting A, only influenza protein families were unknown in the testing set. Additionally, the negative samples in the testing set were generated regardless of human protein families in the PPI pairs. In the Setting B, it is the same as Setting A, but we do not use any human protein families that existed in the training set to generate the negative samples in the testing set, making them inaccessible in the training phase. In the Setting C, both the human and influenza protein families were unknown in the testing set. In the Setting D, following Setting C, we only tested our model on real positive samples. We will have the raw setup as Setting E, allowing the model to train and test PPI data without any constraints. We ended up with 7,614, 7,614, and 1,986 testing PPI pairs under Setting A-C, respectively, with an equal number of positive and negative samples for each. In the Setting D, we obtained 993 positive samples, while in the Setting E, we had 3,178 PPI pairs (1,589 positives and 1,589 negatives).

As shown in **Figure** 3B, we achieved the best performance across all metrics under Setting E for influenza-human PPI prediction, where SEHI-PPI was trained from a comprehensive dataset with a broader range of interaction patterns. Under Setting A, our model also performs well despite the test influenza virus proteins being unknown in the training process. In contrast, Setting B exhibits a slight performance drop (4.5% on average across all metrics) compared to Setting A. This decline likely results from the constraint that host proteins in negative samples are exclusively assigned to either the training or testing dataset, thereby reducing training diversity and limiting generalization capability. When we applied SEHI-PPI under Setting C, where both the influenza and human protein families were entirely novel to the model, we observed a minor-to-moderate decrease in sensitivity, specificity, and MCC. Nevertheless, the model still maintains comparable performance, particularly in the F1 score (0.836) and AUROC (0.837), demonstrating its ability to predict influenza-human PPIs even when neither the interaction pairs nor protein families are previously encountered. To further assess its real-world applicability, we evaluated SEHI-PPI on Setting D, using only experimentally verified positive human-influenza interaction pairs. Remarkably, despite the absence of negative data, SEHI-PPI achieves high accuracy (0.987), precision (1.000), and F1 score (0.993). These results underscore its robustness in reliably identifying true PPIs, even in scenarios where negative interaction samples are unavailable.

#### Comparison of different K-mer segmentation embedding techniques used in SEHI-PPI

We compared the *K*-mer segmentation with different embedding methods, i.e., ProtBERT^50^, DeepWalk^51^, and Node2vec^9^. We used the same experimental setup to train and test the models with these embedding techniques to ensure a fair comparison. The radar chart of **Figure 3**C indicates that ProtBERT consistently outperforms the other two methods across all metrics, achieving superior results in accuracy (0.987), F1-score (0.987), AUROC (0.987), sensitivity (0.986), specificity (0.988), and MCC (0.974). This is probably attributed to the ability of ProtBERT to generate rich, context-aware embeddings that capture intricate biological information of *K*-mer segments. In contrast, Node2vec shows moderate performance with accuracy (0.887), F1-score (0.885), and AUROC (0.886), but struggles with sensitivity (0.771) and MCC (0.794). This result indicates that Node2vec’s structural relationship modeling is helpful but insufficient for capturing deep contextual dependencies. DeepWalk performs the worst, particularly with low sensitivity (0.649) and MCC (0.694), likely due to its reliance on random walk strategies that are less effective for capturing essential sequence dependencies. These results underscore ProtBERT’s capability to extract comprehensive embeddings, enhancing SEHI-PPI’s predictive performance over Node2vec and DeepWalk.

#### Global and local features driving SEHI-PPI performance

To elucidate the contributions of LSTM and GIN modules within the SEHI-PPI model, we conducted ablation experiments by eliminating each module and measuring the impact on model performance. As illustrated in **Figure 3**D, the removal of LSTM leads to a decrease in sensitivity (from 0.986 to 0.888) and MCC (from 0.974 to 0.894), which emphasizes the importance of LSTM in capturing temporal dependencies essential for identifying PPIs. Similarly, excluding the GNN module results in declines in accuracy (from 0.987 to 0.952), specificity (from 0.988 to 0.948), F1-score (from 0.987 to 0.952), and AUROC (from 0.987 to 0.952). These decreases suggest the role of GNNs in providing structural insights into the constructed graphs that enhance the model’s capability to distinguish samples effectively. When using both LSTM and GIN, SEHI-PPI achieves the highest scores across all metrics, demonstrating their synergistic effect. The integration of LSTM and GIN promotes SEHI-PPI to capture comprehensive features, resulting in a robust and well-balanced framework for human-influenza PPI prediction.

### Parameters analysis of SEHI-PPI

#### Effect of K values on the performance of SEHI-PPI

We analyzed the impact of varying *K*-values, i.e., the number of contiguous amino acids in protein sequences within each segmented subsequence, affected SEHI-PPI, specifically tested using the LSTM+GIN architecture. *K*-values were evaluated in the range of 2 to 5, increasing in increments of 1. As shown in **Figure 4**A, the results demonstrate that the SEHI-PPI framework achieves the highest performance metrics when *K*=2, with an accuracy of 0.990 ± 0.002, F1-score of 0.990 ± 0.002, and MCC of 0.980±0.004. As *K* increases to 3, 4, and 5, there is a slight decline in performance across all metrics. At K=3, accuracy, F1-score, and AUROC achieve 0.987 ± 0.002, while further increases to K=4 and K=5 maintain high values, though they are marginally lower compared to K=2.

**Figure 4.**
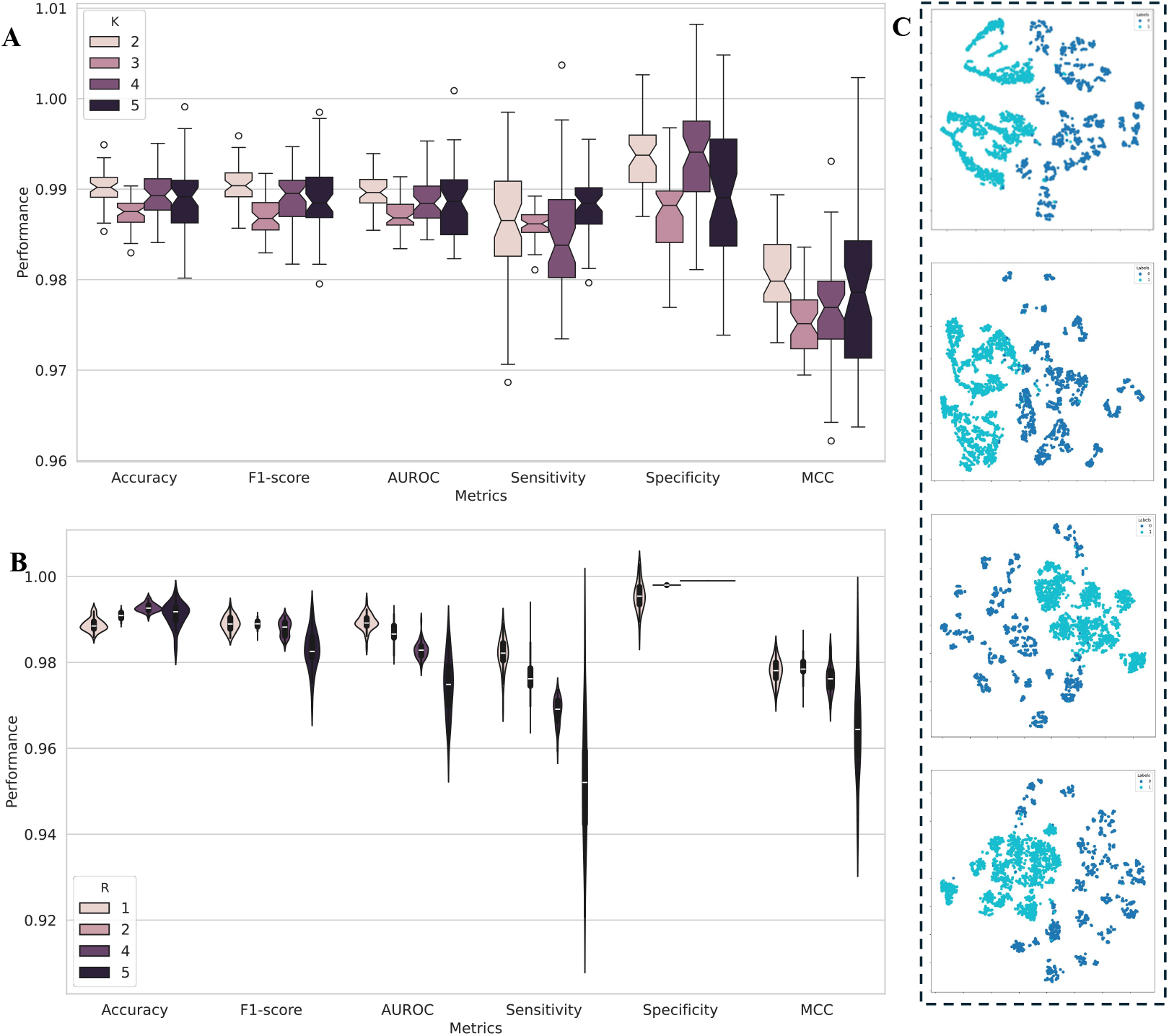
Comprehensive parameter analysis of SEHI-PPI for optimizing performance in human-influenza PPI prediction and graph embedding visualization. (A) The influence of *K* value in *K*-mer on the performance of SEHI-PPI framework (implemented by LSTM + GIN architecture). (B) The influence of R value in positive and negative sample ratio on the performance of SEHI-PPI framework (implemented with LSTM+GIN architecture). (C) Graph embedding visualization of different GNNs in SEHI-PPI model.

#### Effect of R values on SEHI-PPI performance

We further evaluated the effect of the negative-to-positive sample ratio *R* (i.e., negative: positive), ranging from 1 to 5, on the performance of SEHI-PPI implemented with the LSTM+GIN architecture. Figure 4B highlights the variability and distribution of performance metrics as a function of the *R*-values. As *R* increases, the distributions for sensitivity narrow and shift lower, while specificity consistently exhibits minimal variability and remains high. Accuracy, F1-score, and MCC also display a slight downward trend in performance as *R* increases, with the most pronounced variability observed at *R*=5. Particularly, increasing *R* from 1 to 4 leads to improvements in accuracy and specificity, peaking at 0.993 ± 0.001 and 0.999 ± 0.000 respectively when *R*=4. However, this improvement comes with a tradeoff: sensitivity gradually decreases, reflecting the model’s diminished ability to detect positive samples as it becomes more proficient at identifying negative ones. This tradeoff becomes particularly pronounced at *R*=5, where sensitivity drops significantly to 0.951 ± 0.015, even as specificity remains consistently high at 0.999 ± 0.000. F1-score and MCC follow similar trends, with both metrics declining as *R* increases, especially beyond *R*=2. Notably, the most balanced performance is observed at lower *R*-values, particularly at *R*=1 and *R*=2, where both sensitivity and specificity are relatively high, maintaining F1-scores of 0.989 ± 0.002 and 0.989 ± 0.001, respectively, and MCC values of 0.978 ± 0.003 and 0.979 ± 0.003. The slight variance demonstrates that our SEHI-PPI model is robust to the R values, i.e., not sensitive to the ratio of positive and negative samples.

#### Graph embedding visualization of SEHI-PPI with different GNN models

To evaluate the effectiveness of different GNN models in SEHI-PPI, we first extracted the learned graph embeddings, representing protein pair representations as the concatenation of double-view features before the MLP layers were used for final prediction. We then applied t-distributed stochastic neighbor embedding (t-SNE) to visualize these representations in two dimensions, providing insights into the structural differences and clustering patterns across different GNN architectures. As we can see in **Figures 4**C (a) and (b), the positive (interaction, labeled as “1”) and negative samples (non-interaction, labeled as “0”) are partially separated using GAT and PNA models, with most positive samples clustered on the left and negative samples on the right. However, some overlap between the two groups is evident, indicating limited separation capability. In contrast, **Figures 4**C (c) and (d) depict the embeddings learned with the GIN and SGC architectures. These models achieve a clearer separation of positive and negative samples, with distinct clusters and an evident gap between them. These observations indicate that GIN and SGC can better learn the node embeddings compared to GAT and PNA through capturing graph structures, enabling the generation of more meaningful and discriminative node representations, which is consistent with the predictive performance for human-influenza PPIs in **Table 1**.

### Generalization capability of SEHI-PPI on other human-virus PPI predictions

We fine-tuned SEHI-PPI that was trained on the human-influenza PPI dataset by freezing the double-view feature extraction module and learning the new parameters in the fusion layer (i.e., the MLP layer) from other human-virus PPI pairs. We tested our framework on several types of viruses, including Zika virus (Zika, #4,809), Murid herpesvirus 1 (MHV1, #13,160), Human herpesvirus 1 (HHV1, #23,486), Zaire ebolavirus (Ebola, #2,360), severe acute respiratory syndrome coronavirus (SARS-CoV, #12,334), Beta papillomavirus (BPPLM, #6,521), and Human herpesvirus 8 (HHV8, #11,906). We followed the same steps described in the Dataset Construction Module (Method Details) to collect and process human-virus PPI pairs beyond influenza, splitting each species’ dataset into an 8:1:1 ratio for fine-tuning, validation, and testing, respectively. As shown in Table 2, SEHI-PPI demonstrates strong generalization capability, effectively predicting PPIs between humans and other viral species, though with slightly lower performance than human-influenza PPI prediction. This slight decline is likely due to differences in protein sequence composition and functional diversity across viral species, which introduce more complex patterns that challenge the model’s ability to accurately capture global and local features. Nevertheless, SEHI-PPI maintains comparable performance, proving that the learned features were transferable and effective for novel viruses. Notably, our model achieves accuracy, F1-score, and AUROC of 0.960 ± 0.006, with high specificity (0.971 ± 0.016) and sensitivity (0.950 ± 0.014) on the HHV1 dataset, closely approaching its performance on human-influenza PPI prediction. Similarly, high accuracy and superior F1-score and AUROC are observed for MHV1, BPPLM, SARS-CoV, and HHV8. MCC evaluation further confirms a balanced prediction performance across these human-virus PPI pairs. However, the model exhibited slightly lower accuracy on human-Zika (0.882 ± 0.013) and human-Ebola (0.921 ± 0.008) PPI prediction, likely due to smaller dataset sizes, which may have impacted generalization. Despite this, specificity and MCC remain strong, indicating a reliable ability to identify interactive human-virus PPIs, particularly in human-Zika pairs, even with limited data. Overall, SEHI-PPI demonstrates excellent generalizability across diverse viral species, making it a powerful tool for identifying host-virus PPIs in emerging viruses. This capability is crucial for early detection and monitoring of viral threats, as well as guiding the development of targeted therapeutic interventions.

**Table 2.** Evaluation of generalization capability of our SEHI-PPI framework with GIN+LSTM architecture.

| Virus | Accuracy | Sensitivity | Specificity | F1-score | AUROC | MCC |
| --- | --- | --- | --- | --- | --- | --- |
| Zika | $0.882 \pm 0.013$ | $0.906 \pm 0.026$ | $0.858 \pm 0.023$ | $0.882 \pm 0.013$ | $0.882 \pm 0.013$ | $0.765 \pm 0.026$ |
| MHV1 | $0.920 \pm 0.013$ | $0.945 \pm 0.018$ | $0.893 \pm 0.035$ | $0.919 \pm 0.013$ | $0.919 \pm 0.013$ | $0.841 \pm 0.024$ |
| HHV1 | $0.960 \pm 0.006$ | $0.950 \pm 0.014$ | $0.971 \pm 0.016$ | $0.960 \pm 0.006$ | $0.960 \pm 0.006$ | $0.921 \pm 0.011$ |
| Ebola | $0.921 \pm 0.008$ | $0.898 \pm 0.030$ | $0.944 \pm 0.027$ | $0.920 \pm 0.009$ | $0.921 \pm 0.010$ | $0.843 \pm 0.018$ |
| SARS-CoV | $0.934 \pm 0.028$ | $0.909 \pm 0.058$ | $0.958 \pm 0.009$ | $0.934 \pm 0.028$ | $0.934 \pm 0.028$ | $0.870 \pm 0.052$ |
| BPPLM | $0.922 \pm 0.011$ | $0.948 \pm 0.008$ | $0.896 \pm 0.016$ | $0.922 \pm 0.011$ | $0.922 \pm 0.010$ | $0.845 \pm 0.021$ |
| HHV8 | $0.955 \pm 0.007$ | $0.945 \pm 0.014$ | $0.966 \pm 0.027$ | $0.955 \pm 0.007$ | $0.956 \pm 0.007$ | $0.912 \pm 0.014$ |

### Sequence-embedding-based clustering reflects the structural similarities using Alpahfold-3

To further demonstrate the effectiveness of our SEHI-PPI framework and showcase its potential application, we selected two human proteins, i.e., HSPA1B (Heat Shock Protein Family A Member 1B) and RTRAF (RNA Transcription, Translation, and Transport Factor) as examples for case analysis. HSPA1B, known as Heat Shock Protein 70 (HSP70), plays a critical role in cellular stress responses and interacts with viral proteins during influenza virus infection, influencing viral replication and propagation. Using our model, we identified influenza proteins that interact with HSPA1B, including P03485 (M1_I34A1 in Influenza A virus), P03489 (M1_INBLE in Influenza B virus), P03470 (NRAM_I33A0 in Influenza A virus), P03454 (HEMA_I33A0 in Influenza A virus), and P03430 (RDRP_I33A0 in Influenza A virus). To investigate the relationships among these viral proteins, we performed a hierarchical clustering analysis, as illustrated in **Figure 5**A. The results show that M1_INBLE (P03489) and M1_I34A1 (P03485) - matrix protein 1 (M1) from Influenza B virus (strain B/Lee/1940) and Influenza A virus (strain A/Puerto Rico/8/1934 H1N1), respectively-are closely clustered. Despite belonging to different influenza types, their close proximity in the dendrogram suggests substantial sequence or structural similarity, indicating potential functional or evolutionary connections. These findings are further validated through their aligned 3D structures generated by AlphaFold-3 (see Supplementary Data 1), displaying significant overlap across most regions (depicted on the far-left side of **Figure 5**A). Moreover, NRAM_I33A0 (P03470) and HEMA_I33A0 (P03454), corresponding to neuraminidase (NA) and hemagglutinin (HA) proteins respectively, form a distinct cluster. This clustering reflects their complementary roles during the viral lifecycle: HA facilitates viral entry by binding to sialic acid on host cells, while NA cleaves sialic acid to enable viral release^52^. Their close similarity, evident from both their clustering in the dendrogram and the aligned 3D structures (shown on the far-right side of **Figure 5**A), highlights their functional interdependence as critical surface glycoproteins. Overall, these clustering results suggest that the viral proteins interacting with the same host protein may share conserved mechanisms or adaptations. To further analyze human-influenza protein interactions, we visualized both the overall structure (left) and the detailed binding interface (right). The cartoon models show the overall shape and orientation, while the stick models highlight key residues involved in hydrogen bonds and hydrophobic contacts. **Figures 5**C and **5**D show the interactions predicted by the SEHI-PPI model for HSPA1B with influenza proteins M1_I34A1 and M1_INBLE, respectively. The well-defined interaction interface suggests a strong and specific binding, potentially playing a key role in viral replication by ensuring proper M1 localization and stability. Similarly, the interaction between HSPA1B (light pink) and M1_INBLE (green) suggests that HSPA1B may facilitate viral replication. The calculated atomic distances of 3.9 Å and 4.4 Å further confirm strong molecular interactions, reinforcing the stability.

**Figure 5.**
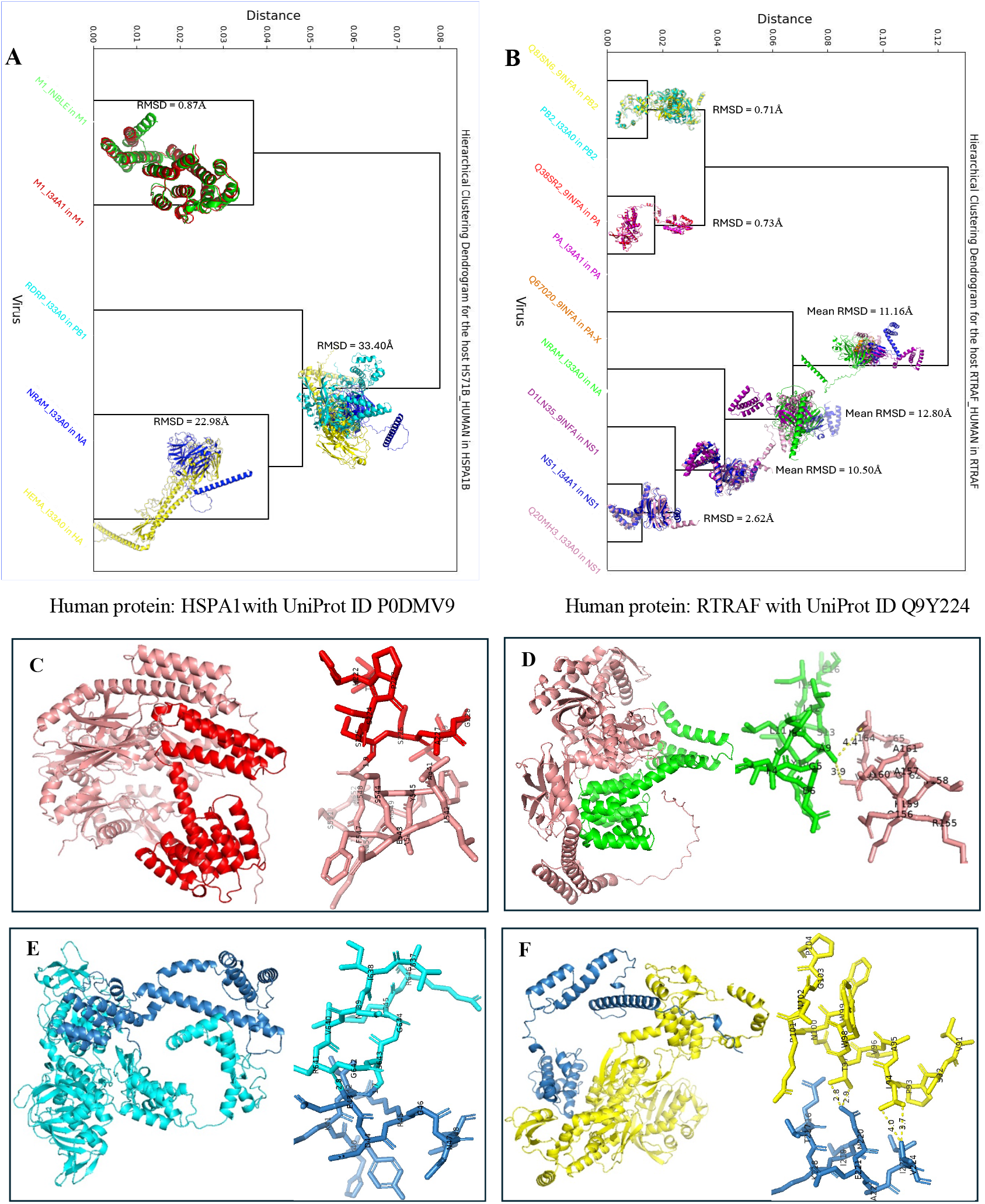
Hierarchical clustering of influenza proteins based on learned embeddings from SEHI-PPI, with structural alignments predicted by AlphaFold-3 and their similarity measures (RMSD), illustrating interactions with two selected target human proteins. (A) Clustering of five influenza proteins interacting with HS71B_HUMAN (HSPA1B): M1_I34A1 (M1, Influenza A), M1_INBLE (M1, Influenza B), NRAM_I33A0 (NA, Influenza A), HEMA_I33A0 (HA, Influenza A), and RDRP_I33A0 (PB1, Influenza A). (B) Clustering of nine influenza proteins interacting with RTRAF_HUMAN (RTRAF): Q38SR2_9INFA (PA, Influenza A), NRAM_I33A0 (NA, Influenza A), NS1_I34A1 (NS1, Influenza A), Q8JSN6_9INFA (PB2, Influenza A), PB2_I33A0 (PB2, Influenza A), Q67020_9INFA (PA-X, Influenza A), Q20MH3_I33A0 (NS1, Influenza A), PA_I34A1 (PA, Influenza A), and D1LN35_9INFA (NS1, Influenza A). (C) and (D) depict the global cartoon and local stick visualizations of HSPA1B interacting with M1_I34A1 and M1_INBLE, while (E) and (F) show similar visualizations for RTRAF interacting with PB2_I33A0 and Q8JSN6_9INFA.

Similarly, RTRAF (RNA Transcription, Translation, and Transport Factor), a human protein integral to RNA metabolism, plays a significant role in human-influenza protein interactions. We also identified several viral proteins predicted to interact with RTRAF using SEHI-PPI, containing Q38SR2 (Q38SR2_9INFA), P03470 (NRAM_I33A0), P03496 (NS1_I34A1), Q8JSN6 (Q8JSN6_9INFA), P03427 (PB2_I33A0), P03433 (PA_I34A1), Q67020 (Q67020_9INFA), D1LN35 (D1LN35_9INFA), and Q20MH3 (Q20MH3_I33A0). These structures predicted by AlphaFold-3 are referred to in Supplementary Data 2. The clustering analysis of these viral proteins, as depicted in **Figure 5**B, uncovers two distinct groups, each reflecting their unique roles in the influenza virus lifecycle. Cluster 1 (the upper part) consists of Q8JSN6, P03427, Q38SR2, and P03433, which collectively form the core transcriptional and replication machinery of the virus. Within this cluster, Q8JSN6 and P03427 (PB2 segment) exhibit high similarity, as both bind to the 7-methylguanosine cap of host pre-mRNAs to initiate the cap-snatching process^53^ (shown on the aligned 3D structures of **Figure 5**B). P03433 (PA segment) and Q38SR2 are grouped due to their shared roles of integral components of the influenza virus heterotrimeric polymerase complex, which orchestrates viral RNA transcription and replication^54^. Cluster 2 (the lower part) involves P03470, P03496, Q67020, D1LN35, and Q20MH3, showcasing distinct yet complementary roles in supporting influenza virus pathogenesis. For example, P03496 (NS1_I34A1) and Q20MH3 form the closest sub-cluster, followed byD1LN35, highlighting their roles in disrupting host RNA processing, inhibiting stress responses, and suppressing antiviral signaling pathways to create a favorable environment for viral replication^55^. NRAM (P03470) contributes by facilitating the release and spread of infectious virions, while Q67020 plays a pivotal role in supporting replication through the regulation of host transcription^56^. **Figures** 5E and 5F illustrate the interactions between RTRAF and the influenza proteins PB2_I33A0 and Q8JSN6_9INFA, respectively. The interaction between RTRAF (light cyan) and PB2_I33A0 (dark blue) is depicted through global and local structural representations, emphasizing their binding interface and overall conformation.

The strong interaction suggests that RTRAF may facilitate viral RNA synthesis or replication by engaging with PB2, a key polymerase subunit essential for influenza transcription. Additionally, the interaction between RTRAF (yellow) and Q8JSN6_9INFA (blue) is visualized through global (left) and local (right) representations, highlighting a stable binding interface.

## Discussion

In this work, we proposed SEHI-PPI, a novel framework for human-influenza PPI prediction, integrating adaptive negative sampling and double-view sequence feature extraction. To construct a high-quality dataset, SEHI-PPI collected large-scale positive human-influenza PPIs from publicly available sources and generated high-quality negative samples using an adaptive negative sampling strategy, ensuring a more informative training process. The model then employed RNNs to capture global sequence dependencies and GNNs to extract local structural features, providing a comprehensive representation of human-influenza PPI interactions. Extensive experiments demonstrated SEHI-PPI’s superior performance over classic classifiers, deep learning models, and LLMs. Notably, our model has shown great capacity for identifying novel human-influenza PPI pairs under different settings (**Figure** 3B). Even when both human and influenza proteins are unknown, it still achieved an F1 score of 0.836 and an AUROC of 0.837. The ablation studies revealed how the different parameters and model architectures could affect the human-influenza PPI prediction. We further evaluated SEHI-PPI’s generalizability by fine-tuning the model and predicting PPI between humans and other viruses, including Zika, MHV1, HHV1, Ebola, SARS-CoV, BPPLM, and HHV8 (**Table** 2). The results have shown that our proposed framework achieved comparable performance when applied to the PPI prediction of other human-virus pairs. We also conducted hierarchical clustering on the influenza proteins identified by SEHI-PPI as binding to the same human protein. The results revealed structural alignments among influenza proteins within the same cluster, as predicted using AlphaFold-3, further proving the effectiveness of our SEHI-PPI framework (**Figure** 5).

Several factors contribute to the superior performance of our SEHI-PPI. First, adaptive negative sampling ensures that the model is trained with high-quality negative samples, making it more robust in distinguishing true interactions from false positives. Unlike random negative sampling, which may introduce easily separable non-interacting pairs, our adaptive approach allows the model to learn a more realistic distribution of interactions, making it particularly effective in predicting novel human-virus PPIs. Second, the double-view feature extraction module strengthens SEHI-PPI’s ability to learn potential biologically meaningful patterns. By combining RNNs for global sequence dependencies and GNNs for local connectivity, the model efficiently captures both long-range functional relationships and localized structural interactions. This dual representation allows SEHI-PPI to generalize better across diverse protein families, which is crucial for accurate PPI prediction. Third, optimized embedding selection and hyperparameter tuning significantly enhance SEHI-PPI’s predictive accuracy. Through extensive benchmarking, our analysis identified ProtBERT embeddings as the most informative sequence representations, outperforming other embedding techniques. These optimizations allow SEHI-PPI to effectively generalize to unseen interactions, making it a versatile and scalable framework, capable of extending beyond human-influenza PPIs to predict interactions in other host-virus systems.

To evaluate SEHI-PPI’s effectiveness, we conducted case studies using two human proteins, HSPA1B and RTRAF, identifying all binding viral proteins using SEHI-PPI. A hierarchical clustering analysis of these viral proteins, combined with AlphaFold-3 structural alignment, revealed that proteins binding to the same human target tend to cluster closely based on structural and functional similarity. A direct observation is that influenza proteins with the same or similar functions are consistently clustered together and exhibit small Root-Mean-Square Deviation (RMSD) values, indicating high structural similarity. For instance, M1_I34A1 and M1_INBLE both belong to the M1 protein family, while Q8JSN6_9INFA and PB2_I33A0 are grouped under the PB2 protein family. This pattern is also evident in other protein clusters, suggesting that SEHI-PPI effectively captures meaningful protein representations, thereby enhancing its ability to accurately predict human-influenza PPIs. Interestingly, when clustering human protein HSPA1B, we observed that despite belonging to different influenza types, influenza proteins exhibited well-aligned 3D structures, suggesting conserved binding mechanisms. For instance, M1_I34A1 and M1_INBLE belong to Influenza A and B respectively, but showed a RMSD of 0.87Å, indicating high structural similarity. Similarly, two NS proteins, NS1_I34A1 and Q20MH3_I33A0, which bind to RTRAF, formed a closely related cluster with an RMSD of 2.62Å, further confirming their structural resemblance. In contrast, other RTRAF-binding viral proteins, such as Q67020_9INFA (PA-X) and NRAM_I33A0 (NA) from Influenza A, displayed greater structural divergence from NS proteins like Q20MH3_I33A0, reflecting their distinct roles in host interaction and viral infection. This clustering analysis further validates SEHI-PPI’s reliability in distinguishing structurally and functionally related viral proteins, reinforcing its potential for broader applications in host-virus interaction research.

While the SEHI-PPI model represents a significant advancement in predicting human-influenza PPIs, several limitations should be acknowledged. (1) *Dependency on high-quality and comprehensive PPI datasets*. Though, to the best of our knowledge, this is the largest human-influenza PPI dataset collected from experimentally validated databases, variations in data quality and completeness across different resources remain a concern. These inconsistencies can lead to noise and impact the prediction performance of our model. (2) *Potential bias from adaptive negative sampling*. Although adaptive negative sampling improves model robustness by generating high-quality negative samples, it may introduce bias if the generated samples fail to represent the true diversity and distribution of non-interaction protein-protein pairs. (3) *Limited interpretability of deep learning framework*. The SEHI-PPI model functions as a “black box”, making it difficult to discern the specific biological mechanisms underlying its predictions. (4) *Reliance on experimental validation*. The study primarily uses computational evaluations to validate the model’s performance. While these results are encouraging, experimental validation of the predicted PPIs is necessary to confirm their true biological function of human-virus protein-protein pairs. Without such confirmation, the reliability and practical applicability of the predictions remain uncertain. There are several avenues for further development of the SEHI-PPI model. First, integrating 3D protein structural data will provide a richer spatial understanding of PPIs, improving predictions beyond sequence-based representations. Advances in AlphaFold-3 make this increasingly feasible, allowing the model to leverage protein conformational changes in interaction prediction. Second, we can incorporate protein mutation analysis and extend the framework to predict protein binding sites and further enhance its applicability. This would enable SEHI-PPI to identify how mutations influence PPIs and help uncover potential therapeutic targets, providing deeper biological insights into protein function and interaction mechanisms. Finally, we will fine-tune and evaluate our proposed framework on other types of PPI, including those relevant to cancer and Alzheimer’s Disease. This would validate its versatility and generalization capabilities, which not only demonstrate SEHI-PPI’s utility in diverse biological contexts but also open new avenues for understanding disease-specific PPIs and guiding targeted therapeutic strategies.

## Methods

Our proposed SEHI-PPI model mainly comprises three components: (1) the dataset construction module; (2) the double-view sequence feature extraction module; and (3) the feature fusion module for human-influenza PPI prediction. We introduce the details in the following subsections.

### Dataset Construction Module

#### Positive Human-Influenza PPI Data Collection

To construct a comprehensive dataset of positive human-influenza protein interaction pairs, we performed a multi-step approach. First, we comprehensively searched available host-virus PPI databases^57^ from public resources, which would contain human-influenza protein-protein interaction pairs experimentally validated. We ended up with several public data sources, including BioGRID^58^, EBI-GOA-nonIntAct^59^, HCVpro^60^, HPIDB^61^, HVIDB^62^, IntAct^63^, VirusMINT^64^, PHISTO^65^, Viruses.STRING^66^, VirHostNet^67^, and VirusMentha^68^. Second, we used the keywords “human” and “influenza” for host and virus types respectively to derive the eligible PPI pairs. We retained only the PPI pairs with valid UniProt IDs for both human and influenza proteins to enable subsequent sequence extraction. Third, we first mapped these UniProt IDs from Swiss-Prot, a high-quality protein database that consists of 572,619 types of manually curated and annotated protein information across a wide range of species. Then, we extracted the corresponding accession codes (AC) and protein sequences of influenza and humans. However, the relatively small number and diversity of proteins in Swiss-Prot limited its coverage. To address this, we supplemented the dataset with sequences from the TrEMBL section of UniProtKB, which offered a broader range of automatically annotated protein sequences, including newly identified ones. Combining Swiss-Prot and TrEMBL data ensured a balance between quality and coverage. Fourth, we further processed the filtered PPI data based on the following steps: i) instances of multiple AC codes associated with the same sequence were consolidated, and duplicate AC entries were removed to create a non-redundant and consistent dataset, ii) missing AC codes, caused by coverage gaps or deleted entries, were identified by comparing the saved AC codes with the expected list, iii) missing sequences were retrieved using online tools such as BioPython, UniProt batch searches, and additional databases to ensure completeness. Lastly, the processed data were saved in CSV format, including four columns: influenza protein AC, human protein AC, influenza protein sequence, and human protein sequence. This process resulted in a high-quality dataset of 15,889 positive human-influenza PPI pairs, encompassing 12 influenza proteins and 4,277 human proteins. Additionally, to evaluate the generalizability of our proposed framework, we also collected interactive protein-protein pairs of other human-virus types (e.g., Zika, Ebola, and SARS-CoV) following the same procedures.

#### Adaptive Negative Human-Influenza PPI Data Generation

Random negative data sampling^35,69^ is widely adopted for generating negative samples in PPI prediction due to its simplicity and ease of implementation. It involved selecting negative samples from unobserved human-influenza pairs, but this approach may comprise data quality issue^70^. To address this problem, we proposed an adaptive negative sampling strategy that leverages sequence similarities between human and influenza sequences to improve the quality and diversity of negative samples. As illustrated in Supplementary Fig. 1, we assumed a positive human-influenza PPI pair was represented as (*v, h*), where 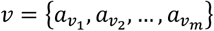 and 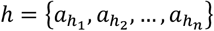 were influenza and human sequence respectively, consisting of *m* and *n* amino acids. We started with building a candidate set 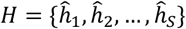 for a given influenza sequence *v* by selecting human sequence 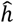 that has never been observed to interact with *v*, ensuring 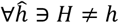. Notably, a *K*-mer is a subsequence of length *K* that is derived from a longer biological sequence, such as a DNA, RNA, or protein sequence. It is commonly used in bioinformatics to represent sequence information in a fragmented form, enabling efficient computational analysis. Both the positive host sequence *h* and candidate negative sequences 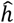 were then segmented into overlapping *K*-mer sub-sequences (segmentations), such as 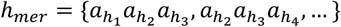 and 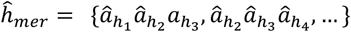, respectively, where *K*=3 in this example. The sequence similarity between the *K*-mer segmentations of the sequences was then computed based on Eqn. (1) below:

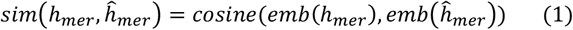

where *emb*() is a function to get the average embeddings of *K*-mer segmentations, implemented by the pretrained protein language models (here we used ProtBERT^50^) and *consine*() is the cosine similarity function. We then ranked all host sequences in the candidate set *H* in a descending way based on the sequence similarities to get the sorted set 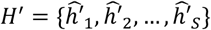. Next, we introduced a ranking position generator to further decide which negative sample should be selected based on the sorted set *H′* following Eqn (2):

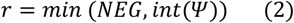

where *NEG* is a constant to indicate the maximized number of negative samples, and Ψ is generated from a predefined distribution as is shown in Eqn (3):

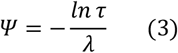

where τ is a random value from Gaussian distribution and λ is the rate value. When τ is set within the range (0,1] with a Gaussian distribution and λ = 1, Ψ is typically a small value. This implies that *r* is highly likely to be a small integer. Consequently, this can lead to a large gradient magnitude during the training of a deep learning model, potentially affecting the stability and convergence of the training process. Finally, we selected the host sequence ranked at position *r* from the ordered candidate set 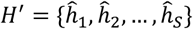 to form the negative influenza and human sequence pair denoted as 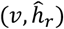. By performing this adaptive negative sampling strategy, we can (1) obtain the host sequences with relatively larger sequence similarity scores which are more likely to be selected; (2) explore the host sequences that have never been interacted by the influenza sequence for promoting diversity.

#### Sequential and Graph Structures Creation

After constructing the positive and negative human-influenza PPI pairs, we segmented them into overlapping *K*-mer sub-sequences. To fully utilize the ability of pre-trained protein language models (PLMs) for processing protein sequences, we used them to initialize the embedding of *K*-mer sub-sequences. In this study, we employed ProtBert^50^, a transformer-based pre-trained model designed specifically for protein sequence and function. The segmented sub-sequences were connected based on the orderliness of sequential data, i.e., (*v*_*mer*_, *h*_*mer*_). To address the variability in sequence lengths of human and influenza sequences, we aligned all sequences to a predefined fixed length, denoted as *Max*, to ensure compatibility with RNN models. This alignment enabled all proteins to be represented consistently, regardless of their original lengths, facilitating uniform encoding while preserving meaningful sequence information up to the specified length. Particularly, if the length of the sequence was smaller than the defined maximum length, they would be padded with “0” at the right hand of the sequence. We further constructed de Bruijn graphs^71^ to represent these sub-sequences, i.e., *v*_*mer*_ and *h*_*mer*_, where each sub-sequence was a node within the graph. A de Bruijn graph is a data structure that presents overlaps between sequences, with nodes as *K*-mer and edges indicating *K*-1 overlaps. We constructed these *K*-mers as nodes and built edges according to the order of *v*_*mer*_ and *h*_*mer*_ to create graphs, i.e., *G*^*v*^ and *G*^*h*^ with the node features 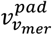 and 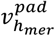. To enhance the importance of edges (i.e., *e*_*ij*_), we then assigned weights to each directed edge, with each weight representing the frequency of an edge that connected two nodes in the graphs. Further, to mitigate the impact of the absolute difference between edge frequencies, we normalized the edge weights in the graph with Eqn. (4):

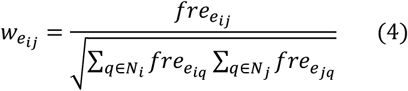

where 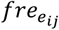 denotes the frequency weight of the edge from node *i* to node *j*, and *N*_*i*_ is the set of neighbor nodes of node *i*. Finally, we got the weighted directed graphs of influenza and human, denoted as 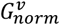 and 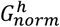 with node features 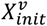 and 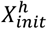, and edge weights 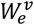 and 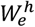, respectively.

### Double-view Feature Extraction Module

Notably, traditional sequence-based methods^27,70^ for human-influenza PPI prediction, such as feature engineering techniques (e.g., Conjoint Triad, Local Descriptor) or models like CNNs, and GNNs, often failed to capture both global and local sequence features simultaneously. Here, we proposed a double-view feature extraction module that combined RNNs and GNNs to extract the global long-term dependencies and local features from input sequences, respectively. This integration enabled the model to comprehensively capture both overarching sequence patterns, thereby enhancing the accuracy and robustness of human-influenza PPI predictions.

#### Global-view Sequence Feature Extraction

Global-view features of protein sequences represented the comprehensive patterns and characteristics of the entire sequence, providing insights into the protein’s overall function and role in biological processes. This helped to capture long-range dependencies and enabled tasks requiring an understanding of the entire sequence. Given the padded *K*-mer segmentations of influenza and human proteins sequences with the initial embeddings, denoted as 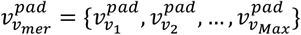 and 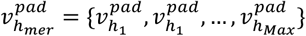, we utilized LSTM to capture the global features for influenza and human protein sequences with Eqn. (5).

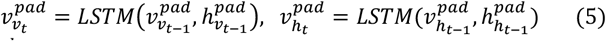

where 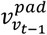 and 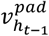 represent the *K*-mer segmentation embeddings of influenza and human proteins at the step *t* − 1, 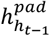 and 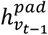 denote the hidden states of influenza and human proteins at the step *t* − 1(1 ≤ *t* ≤ *Max*). To explore the interaction between influenza and human proteins, we concatenated the hidden state of the step *Max* for the influenza and human, i.e., 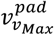 and 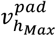, respectively to capture the global-view features, denoted as 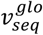. Another type of sequence encoding method, GRU (Gated Recurrent Unit), was also utilized, with detailed information provided in Supplementary Note 1.

#### Local-view Sequence Feature Extraction

Local-view features focused on specific short-range sequence elements critical for precise interactions, emphasizing regions essential for functional activities and interactions for specific protein behaviors. To capture the comprehensive sequence features, we utilized GNN models to capture the local-view features, and here we employed graph isomorphism network (GIN)^72^ as the example to illustrate the process. Other types of GNN models in the experiments can be found in Supplementary Note 1. Generally, GNN models learn node embeddings through the aggregate function and update function. The aggregate function focused on aggregating information from neighbors while the update function aimed at updating the information of the target node from the current layer and previous layer. The process was shown with Eqn. (6):

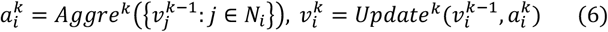

where *N*_*i*_ is the neighbor of node *i*. The main idea lies in message passing and aggregation mechanisms that iteratively update node representations by considering the local neighborhood structure while being invariant to node permutations. The node representations were updated with Eqn. (7):

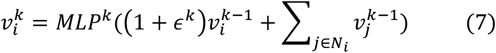

where *N*_*i*_ represents the neighborhood of node *i* and ϵ^*k*^ is a parameter to preserve permutation invariance at the *k*-th iteration. MLP, short for multi-layer perceptron, is used to transform aggregated information by combining the current node’s representation with the sum of its neighbors’ representations. This transformation is achieved through the integration of an aggregate function, which gathers information from neighboring nodes, and an update function, which refines the node’s representation. We then employed GIN architecture, denoted as *GINConv*(), to learn the node embeddings given the graphs of influenza and human proteins, i.e., 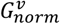 and 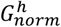 owning initial node features 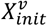 and 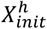, and edge weights 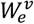 and 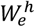, where the updated node features were shown with Eqn. (8):

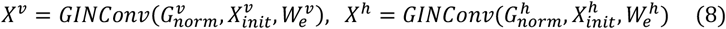

Subsequently, the graph embeddings of influenza and human proteins are captured by the pooling strategies, such as mean pooling, max pooling, and sum pooling, with Eqn. (9).

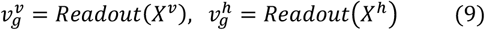

where *Readout*() is the pooling function. To explore the interaction between influenza and human proteins at the local view, we concatenate the learned graph embeddings for the local-view features, denoted as 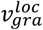.

### Global and Local Features Fusion Module

After extracting global-view and local-view sequence features, we fused them to obtain comprehensive information about human and influenza proteins. This was achieved by concatenating the global-view sequence features 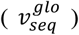 with the local-view graph features 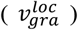, represented as 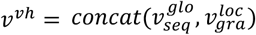, where *concat*() denotes the concatenation operation. The fused feature representation *v*^*vh*^ was then input into Multi-Layer Perceptron (MLP) layers, which served as a classifier to predict the probability of interaction that the influenza *v* and the human *h* interact with each other with Eqn. (10).

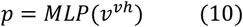

### Model Training

If the influenza *v* and the human *h* interact, the label is 1, otherwise 0. Then, we treated the human-influenza interaction problem as the binary classification task. The objective function was shown with Eqn. (11):

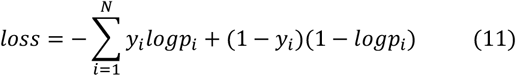

where *y*_*i*_ is the ground truth label and *p*_*i*_ is the predicted label. *N* is the number of samples.

### Experimental Setup

#### Baseline Models

We compared our proposed SEHI-PPI model with several groups of baseline models, including (1) traditional machine learning (ML) methods, (2) existing deep learning (DL) methods, and (3) Large Language Model-based methods. Specifically, in ML methods, we implemented Gaussian Naïve Bayes (GNB), logic regression (LR), support vector machine (SVM), decision tree (DT), K-nearest neighbors (KNN), XGBoost (XGB), and random forest (RF) as the classifiers where the averaged *K*-mer segmentation embeddings were used as the features to feed these classifiers. In DL methods, we compared CNNLSTM, HGCN, DSSGNN, and HIGH. CNNLSTM was a hybrid deep learning framework that integrated four convolutional neural network (CNN) layers with a long short-term memory (LSTM) network to predict human-influenza protein-protein interactions (PPIs) using only protein sequence information^42^. Hyperbolic graph convolutions network (HGCN) was a generalization of inductive GCNs in hyperbolic geometry from Euclidean space to hyperbolic geometric space to obtain smaller distortion embeddings, to predict PPI with the sequence data^43^. DSSGNN (dual-level graph neural network) was a sophisticated model for multi-categorical PPI prediction, which employed a dual-level architecture: a protein-level GNN to extract features from individual proteins and an interaction-level GNN to predict specific interaction categories between protein pairs^44^. HIGH was a hierarchical graph learning model for predicting PPIs and understanding their molecular mechanisms, which represented PPIs in a two-tier graph structure: a macro-level PPI network and a micro-level protein graph to capture structural and chemical features of individual proteins^45^. In LLM baselines, we leveraged two versions of GPT and LLAMA with different numbers of shots, e.g., 0, 2, and 4 for comparison. The shot number here refers to the quantity of labeled data consumed by LLMs. GPT-3.5 was an advanced language model developed by OpenAI that enhanced the capabilities of its predecessor, GPT-3, by providing more coherent and contextually relevant responses^46^. GPT-4 built upon the strengths of GPT-3.5, introducing significant advancements in reasoning, problem-solving, and comprehension of complex instructions^47^. LLAMA2, developed by Meta AI, was an open-source large language model that delivers high performance in natural language processing tasks while promoting transparency and accessibility^48^. LLAMA3 continued the trajectory set by LLAMA2, offering improved efficiency, scalability, and accuracy in processing language tasks^49^. The used resources are shown in Supplementary Table 3.

#### Implementation Details

The SEHI-PPI model was implemented with PyTorch and trained on a single A100 GPU. Human-influenza protein sequence pairs were labeled as “1” for positive interactions and “0” for negative interactions, forming a balanced dataset of 31,778 samples with a 1:1 ratio of positive to negative pairs. The dataset was split into training, validation, and testing sets in an 8:1:1 ratio. For double-view feature extraction, the global-view sequence features were encoded using a two-layer LSTM, while the local-view sequence features were encoded using a two-layer GIN. The classifier consisted of two MLP layers. The model was trained with a batch size of 32 and a learning rate of 0.001, optimized using Adam. Overfitting was mitigated with a dropout strategy (rate of 0.25) and early stopping, which halted training if performance did not improve for 20 consecutive epochs. The training process spanned 80 epochs and included 5-fold cross-validation with different random seeds. Evaluation metrics were reported as the mean and standard deviation across folds, ensuring robust performance assessment.

#### Evaluation Metrics

The performance of the SEHI-PPI model and baseline methods was evaluated on the testing set using six metrics: Accuracy (ACC) to measure overall correctness, F1-score (F1) to balance precision and recall, Area Under the Receiver Operating Characteristic Curve (AUROC) to assess discriminative ability, Sensitivity (SN) to evaluate true positive rate, Specificity (SP) to measure true negative rate, and Matthews Correlation Coefficient (MCC) to capture the overall quality of binary classifications, accounting for imbalances in the dataset.

## Data availability

The PPI data used in this study includes BioGRID, EBI-GOA-nonIntAct, HCVpro, HPIDB, HVIDB, IntAct, VirusMINT, PHISTO, Viruses.STRING, VirHostNet, and VirusMentha (https://github.com/UF-HOBI-Yin-Lab/SEHI-PPI/blob/main/Supplementary%20Material/interactiondata.pdf). Protein sequence data is available in UniProt database (https://www.uniprot.org/). The protein structures are obtained by AlphaFold-3: https://alphafoldserver.com/. All other relevant data supporting the key findings of this study are available within the article and its Supplementary Information files or from the corresponding author upon reasonable request. Source data are provided with this paper.

## Code availability

All codes used in this study are available at https://github.com/UF-HOBI-Yin-Lab/SEHI-PPI.

## ACKNOWLEDGMENTS

This study was partially supported by grants from Centers for Disease Control and Prevention (1U18DP006512) and The Elsa U. Pardee Foundation.

## Author information

Authors and Affiliations

**Department of Health Outcomes & Biomedical Informatics, University of Florida, 1889 Museum Rd, Gainesville, FL, USA**

Qiang Yang & Rui Yin

**Department of Biomedical Engineering, University of Florida, 1275 Center Dr, Gainesville, FL, USA**

Xiao Fan

**Department of Biochemistry & Molecular Biology, University of Texas Medical Branch, 301 University Blvd, Galveston, TX, USA**

Haiqing Zhao

**Department of Molecular Genetics and Microbiology, University of Florida, 1200 Newell Drive, Gainesville, FL, USA**

Zhe Ma & Megan Stanifer

School of Medicine, Indiana University, 340 W 10th St, Indianapolis, IN, USA

Jiang Bian

**Department of Pathology, Immunology and Laboratory Medicine, University of Florida, 1600 SW Archer Rd, Gainesville, FL, USA**

Marco Salemi

### Contributions

R.Y. and Q. Y. conceived the study. Q.Y. performed experiments and data analysis. R.Y., Q.Y., F. X., and H.Z. interpreted the data analysis. Q.Y. and R.Y. drafted the manuscript. Q.Y., R.Y., F. X., and H.Z. critically revised the manuscript. Z.M., M.S., J.B., and M.S. provided suggestions on experiments and manuscript revision. All authors critically revised and gave final approval of the manuscript.

### Corresponding author

Correspondence to Rui Yin.

## Ethics declarations

Competing interests

The authors declare no competing interests.

## Supplementary information

**Supplementary Information**

**Description of Additional Supplementary Files**

**Supplementary Data 1**

**Supplementary Data 2**

## Notes

### Competing Interest Statement

The authors have declared no competing interest.

### Summary of Updates

Introduction section was updated to have a clear structure.

