## Supplementary material for "SEHI-PPI: An End-to-End Sampling-Enhanced Human-Influenza Protein-Protein Interaction Prediction Framework with Double-View Learning": https://github.com/UF-HOBI-Yin-Lab/SEHI-PPI/supp.pdf (from another version of this preprint: version 1, 2025-03-12)

### Supplementary Figures

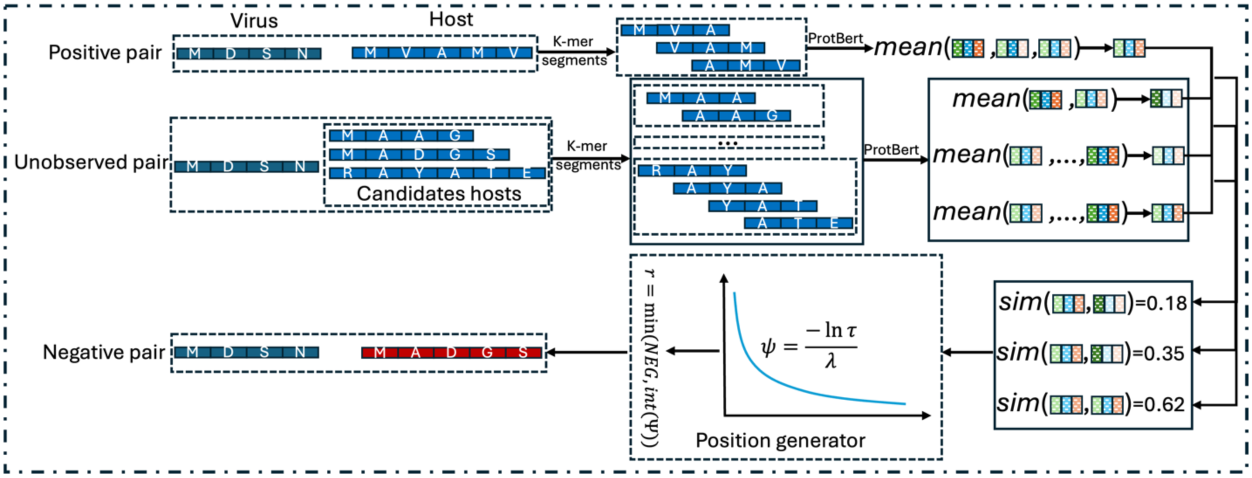


**Supplementary Figure 1**. **The process of the adaptive negative sampling strategy.** Given the positive human-virus sequence pair, it first generates candidate hosts from the unobserved interacted pairs for the target virus sequence. It then generates the overlapping $K$-mer segmentations and calculated the similarities between positive host and candidate hosts. Next, a position generator deciding which candidate was designed to select the negative sample based on the sorted similarities.

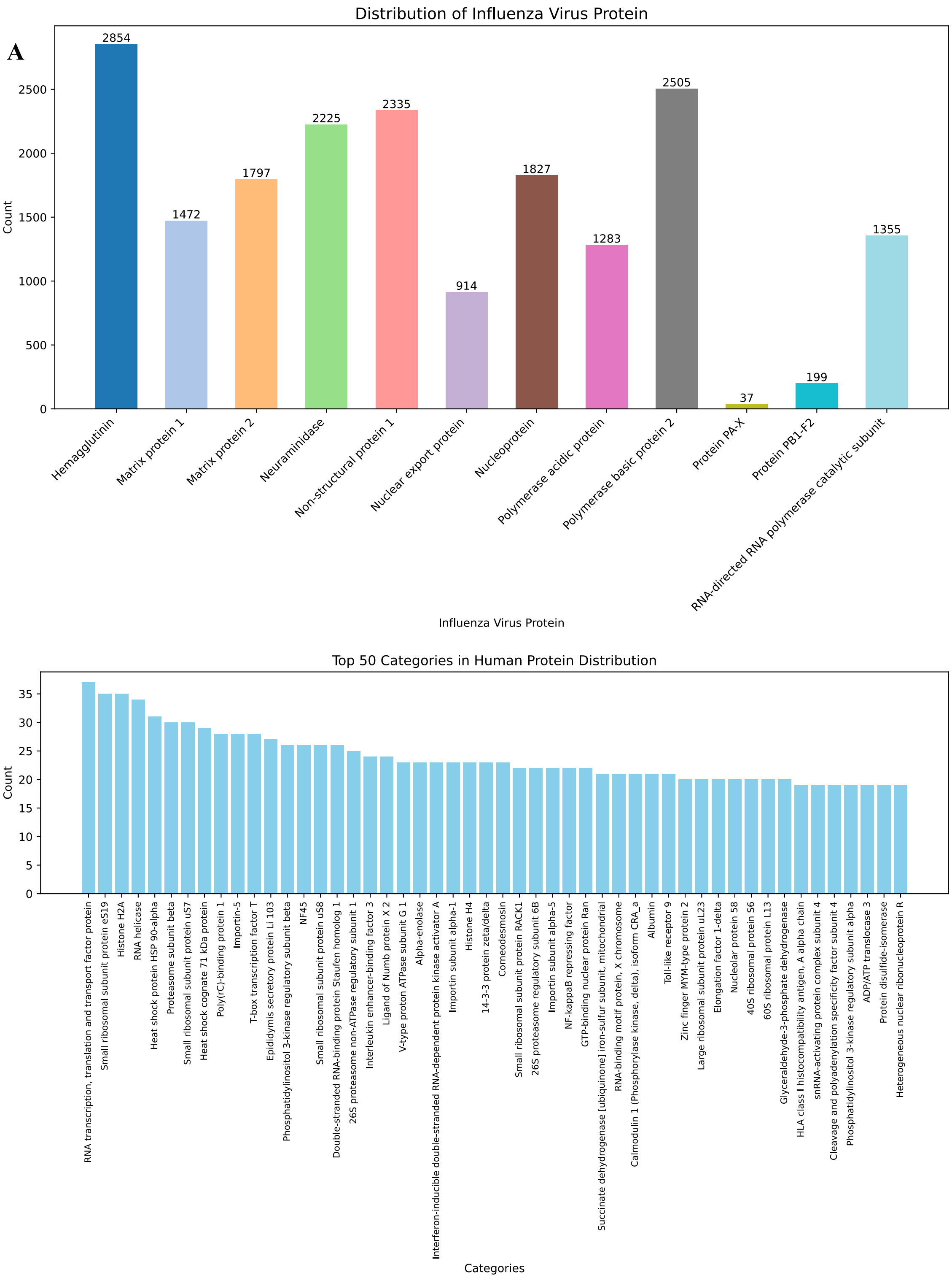


**Supplementary Figure 2. The data distribution of human and influenza protein distribution derived from the collected positive human-influenza PPI pairs.** (A) The statistical information of 12 influenza virus protein types. (B) The statistical information of top 50 in 4277 human protein types.

### Supplementary Tables

**Supplementary Table 1**. Statistics information of the collected positive human-influenza PPI data.

| **Category** | **Count/Range** |
|---|---|
| Human-Influenza Interaction Pairs | 15,889 |
| Unique Influenza Sequence Number | 154 |
| Influenza Protein Type Number | 12 |
| Unique Human Sequence Number | 4,677 |
| Human Protein Type Number | 4,277 |
| Influenza Protein Sequence Length | 18-2,335 (avg: 494) |
| Human Protein Sequence Length | 17-2,335 (avg: 847) |

**Supplementary Table 2. Prompts of in-context learning in LLMs for human-influenza PPI prediction.**

| **Prompt for zero-shot Learning** | **Prompt for few-shot Learning** |
|---|---|
| Task Prompt: Labeling Sequences for Virus-Host Protein-Protein Interaction Prediction<br>Introduction: Welcome to the virus-host protein-protein interaction (PPI) labeling task! Your objective is to determine whether two provided amino acid sequences suggest an interaction between a viral protein (marked as '@VIRUS_PROTEIN$') and a host protein (marked as '@HOST_PROTEIN$'). These sequences are consisted of 20 different types of amino acids. Your task is to label these sequences as either positive (1) for interaction or negative (0) for no interaction, based on the predicted relationship between the viral and host proteins.<br><br>Viral Protein Sequence: @VIRUS_PROTEIN$: {virus_protein_sequence}<br>Host Protein Sequence: @HOST_PROTEIN$: {host_protein_sequence}<br><br>Please output your decision (1 or 0) only.<br>Answer: | Task Prompt: Labeling Sequences for Virus-Host Protein-Protein Interaction Prediction<br>Introduction: Welcome to the virus-host protein-protein interaction (PPI) labeling task! Your objective is to determine whether two provided amino acid sequences suggest an interaction between a viral protein (marked as '@VIRUS_PROTEIN$') and a host protein (marked as '@HOST_PROTEIN$'). These sequences are consisted of 20 different types of amino acids. Your task is to label these sequences as either positive (1) for interaction or negative (0) for no interaction, based on the predicted relationship between the viral and host proteins.<br>{few_shot_exams}<br><br>Now, please analyze the following sequences:<br>Viral Protein Sequence: @VIRUS_PROTEIN$: {virus_protein_sequence}<br>Host Protein Sequence: @HOST_PROTEIN$: {host_protein_sequence}<br><br>Please output your decision (1 or 0) only.<br>Answer: |

**Supplementary Table 3**. **Used resources of SEHI-PPI.**

| REAGENT or RESOURCE | SOURCE | IDENTIFIER |
|---|---|---|
| Deposited data | | |
| UniProtKB database | Coudert et al. | https://www.uniprot.org/ |
| Alphafold3 | Abramson et al. | https://alphafoldserver.com/ |
| Software and algorithms | | |
| Machine Learning classifiers and clustering | https://scikit-learn.org/stable/supervised_learning.html | Version 1.5.0 |
| DeepWalk | Bryan et al. | https://github.com/phanein/deepwalk |
| Node2vec | Aditya et al. | https://github.com/aditya-grover/node2vec |
| ProtBert | Ahmed et al. | https://github.com/agemagician/ProtTrans |
| CNNLSTM | Deng et al. | https://ieeexplore.ieee.org/document/9313117 |
| HGCN | Chami et al. | https://github.com/HazyResearch/hgcn |
| DSSGNN | Chang et al. | https://github.com/cstudy1/DSSGNN-PPI |
| HIGH | Gao et al. | https://github.com/zqgao22/HIGH-PPI |
| GPT3.5 | Brown et al. | https://arxiv.org/abs/2005.14165 |
| GPT4.0 | OpenAI et al. | https://cdn.openai.com/papers/gpt-4.pdf |
| LLAMA2 | Hugo et al. | https://arxiv.org/abs/2307.09288 |
| LLAMA3 | Abhimanyu et al. | https://arxiv.org/abs/2407.21783 |

### Supplementary Notes

### Supplementary Note 1: Variants of deep learning methods in SEHI-PPI

In SEHI-PPI, we leveraged diverse deep learning methods for feature extraction at both global and local views. For global-view feature extraction, we utilized RNNs, specifically the gated recurrent unit (GRU) and LSTM, to capture sequential dependencies in the data. For local-view feature extraction, we employed GNNs, exploring various architectures such as GraphSAGE (SAGE), graph convolution network (GCN), graph attention network (GAT), principal neighborhood aggregation network (PNA), and simple graph convolution network (SGC) to effectively capture structural and neighborhood information within the sequences.

GRU, introduced by Kyunghyun Cho et al., is a type of RNN architecture optimized for sequential data processing[1]. It simplifies LSTM architecture by utilizing only two gates—the reset gate and the update gate. The reset gate determines the extent to which past information is forgotten, while the update gate controls how much of the new information is retained. This design reduces computational complexity while maintaining effective learning of long-term dependencies in sequences. The relevant equations for GRU are shown below:

$$r_t = \sigma(W_r x_t + U_r h_{t-1})$$

$$z_t = \sigma(W_z x_t + U_z h_{t-1})$$

$$\tilde{h}_t = tanh(W x_t + r_t \odot U h_{t-1})$$

$$h_t = (1 - z_t) \odot h_{t-1} + z_t \odot \tilde{h}_t)$$

where $r_t$ and $z_t$ are the reset and update gates, respectively, and $h_t$ is the hidden state at time *t*. By controlling the amount of past information forgotten and new information retained, GRU effectively updates the hidden states while managing information flow in sequential tasks.

SAGE, introduced by Hamilton et al., is a scalable inductive framework for generating node embeddings in large graphs[2]. It efficiently handles unseen nodes by sampling and aggregating features from local neighborhoods. The propagation rule in SAGE to update the network parameters is defined as:

$$h_i^k = \sigma(W^k \cdot Aggre^k(\{h_j^{k-1}, \forall j \in N_i\}))$$

where $N_i$ represents the neighbors of node $I$, $W^k$ is the learnable weight matrix, and $\sigma$ is a non-linear activation function. The aggregation function such as mean, pooling, or LSTM-based, consolidates features from neighboring nodes. This approach generates a new embedding matrix for each node by incorporating features from its local neighborhood while maintaining scalability.

GCN, proposed by Kipf and Welling, is designed for semi-supervised learning on graph-structured data by employing an efficient adaptation of convolutional neural networks[3]. It performs localized convolution operations by aggregating feature information from a node's neighbors and utilizes spectral graph theory to define convolutional filters. The propagation rule of GCN is formulated by the following equation to update the network parameters:

$$h_i^k = \sigma(\widetilde{D}^{-1/2}\tilde{A}\widetilde{D}^{-1/2}h_i^{k-1}W^{k-1})$$

where $\tilde{A} = A + I$ is the adjacency matrix with self-loops, $\widetilde{D}$ is the degree matrix of $\tilde{A}$, $h_i^{k-1}$ represents the node embeddings of $I$ at layer *k-1*, and $W^{k-1}$ is the weight matrix. The GCN layer aggregated features from neighboring nodes by utilizing normalized edge weights, enabling it to update the embedding matrix. This process effectively combines each node's intrinsic characteristics with the features of its local neighbors, resulting in enriched and context-aware node representations.

GAT, introduced by Veličković et al., incorporates attention mechanisms into GNNs, allowing nodes to assign different weights to their neighbors during feature aggregation[4]. This selective aggregation enables nodes to prioritize specific neighbors or edges, guided by learned attention coefficients, thereby enhancing the model's ability to focus on the most relevant relationships and features. The mathematical equation representing the attention mechanism in GAT is defined as follows:

$$e_{ij} = LeakReLU\left(a^T[W \cdot h_i \parallel W \cdot h_j]\right)$$

where $e_{ij}$ denotes the unnormalized attention coefficient for node $j$ in node $i$'s neighborhood, $a$ is the learnable attention weight vector, $W$ is the transformation weight, and $\parallel$ indicates concatenation. $LeakReLU$ is an activation function that introduces small gradients for negative inputs. After $\alpha_{ij}$ is computed for all node pairs, we normalize the attention coefficients through the SoftMax function on the neighborhood of node $i$.

$$\alpha_{ij} = \frac{\exp(e_{ij})}{\sum_{k \in N_i} \exp(e_{ik})}$$

where $N_i$ is the neighborhood of node $i$, and $exp$ is the exponential function. This allows GAT to dynamically focus on more relevant neighboring nodes, enhancing the relevance of aggregated features.

PNA, proposed by Corso et al., is a GNN architecture designed to effectively capture diverse neighborhood statistics by utilizing multiple aggregators, such as mean, max, and sum, which are combined through a learnable aggregation mechanism[5]. To enhance adaptability across varying graph structures, PNA also integrates degree-scalers, which accounts for differences in node neighborhood sizes, ensuring robust and flexible performance in diverse graph environments. The propagation rule for PNA at layer *k* was given by:

$$h_i^k = \sigma\Big(\sum_{m \in M} \phi_m \left(Aggre_m\left(\left\{h_j^{k-1}, \forall j \in N_i\right\}\right)\right) W_m^k\Big)$$

where $M$ is the set of aggregation functions (e.g., mean, max, min), $\phi_m$ is degree-scalers that adjust for neighborhood size variability, and $W_m^k$ is the weight matrix of the aggregation function of *m* at layer *k*. By combining multiple aggregation methods, PNA effectively represents diverse graph structures, offering a flexible embedding matrix for node *i* through combined neighborhood features.

SGC, proposed by Wu et al., is a streamlined variant of traditional GCNs that removes nonlinear activation functions and collapses weight matrices between layers[6]. By focusing solely on linear transformations and multi-hop neighborhood aggregation, SGC significantly reduces computational complexity while retaining the ability to effectively capture structural information from graphs. The propagation rule of SGC is shown below:

$$H^k = \tilde{A}^k X W$$

where $\tilde{A} = \widetilde{D}^{-1/2} A \widetilde{D}^{-1/2}$ is the normalized adjacency matrix, with $\widetilde{D}$ as its degree matrix, $X$ is the input feature matrix, $W$ is the weight matrix, and $k$ denotes the number of hops. This streamlined approach focuses on aggregating multi-hop neighborhood information with fewer computations, significantly reducing computational complexity and simplifying the training process.

### Supplementary Note 2: Random negative sampling and statistics negative sampling for negative data

Random Negative Sampling (RNS) randomly selects unobserved human-influenza pairs in the dataset as the negative samples. The below equation demonstrates the negative sample generation process:

$$RNS(N) = Rand((v_i, h_j) | (v_i, h_j) \notin P, (v_i, h_j) \in U)$$

where $P$ and U are the observed positive and all possible virus-host pairs respectively, and $Rand()$ is the function to randomly select negative samples from data.

Statistics Negative Sampling (SNS) first analyzes the statistics information of unobserved virus-host pairs using the *K*-mer segmentations and then selects the host sequence with the lowest overlaps for the positive host sequence as the negative samples. Let the overlap between $K$-mer profiles of two sequences $v_i$ and $h_j$ be denoted as $Overlap(v_i, h_j)$. Then, the below equation shows the negative sample generation process:

$$SNS(N) = \{(v_i, h_j) | argmin_{h_j \in U \setminus P} Overlap(v_p, h_j), v_p \in P\}$$

where $Overlap(v_p, h_j)$ represents the statistical similarity (e.g., the same $K$-mer segmentations) between $v_p$ and $h_j$.
